# How simple cell to cell communication rules can generate and maintain scale invariant gradients of signalling activity across a multicellular population

**DOI:** 10.1101/2020.11.03.366633

**Authors:** Jack Oldham

## Abstract

This paper shows computationally and conceptually how gradients of signalling activity can be generated and dynamically maintained across a population of cells using very simple cell to cell communication rules. The rules work on the basis of cells regulating their production rate of a signalling molecule according to the production rates of their immediate neighbours. Highly stable, scale invariant signalling gradients can be formed across the population, with highest rates at the centre and lowest at the periphery.

The cell to cell communication behaviour that causes gradient formation is first explained in a descriptive, thought experiment type manner. It is then defined more formally using a conceptual, mathematically discrete computational model, which provides a network or graph type framework in which it is easy to analyse and control discrete signals that are sent between neighbouring cells. This provides an intuitive method of explaining how the signalling gradient emerges as a result of local cell to cell communication. Finally, examples of gradient formation are shown using software implementations of the model.

## Introduction

The concepts in this paper are derived from a Quorum Sensing (Miller & Bassler 2001) inspired thought experiment that is to ask how a population of cells can collectively and synchronously detect their population size using only local cell to cell communication. The algorithm used to achieve this consequently results in formation of a gradient of signal production rates across the population of cells (fig 1). This gradient forming behaviour is arguably interesting because, in a conceptual sense at least, it has some resemblance to morphogen gradients observed in real biological systems (Blair 2007, Hill 2017, O’Connor et al 2006, Placzek & Briscoe 2018, Wolpert 1969). Whilst there are probably many ways to conjure up a mechanism to form some kind of signalling gradient using local signalling rules, this seemed notable due to the simplicity of the algorithm and the fact that it is achieved primarily through use of autoregulation, which is a commonly observed characteristic of morphogen signalling in developing organisms (Ashe and Briscoe 2006, Crews & Pearson 2009, Schier 2009, Zecca & Struhl 2007).

**Figure 1 -.**
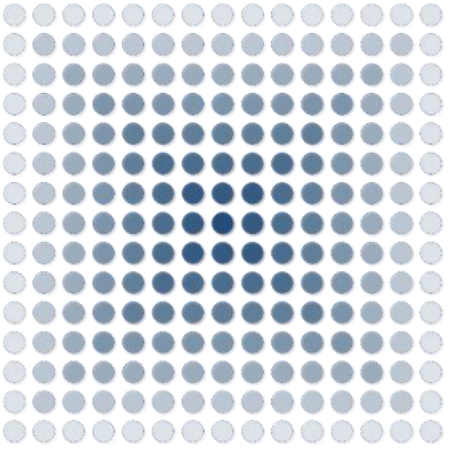
Conceptual diagram of a gradient of signalling activity generated by a population of cells. The strength of blue indicates the rate of signalling of each cell; the stronger the blue, the faster the rate at which that cell is emitting signalling molecules. Signalling molecules themselves are not shown.

Even though there is a conceptual reference being made to morphogen gradients, and to some extent quorum sensing, this paper is not in any way making a claim or hypothesis that real biological systems use the same or similar mechanisms to those described here. This is because the model used is highly conceptual and there is also no empirical evidence directly linking any of the ideas in this paper to real biological systems. Therefore, the aim of this paper is only to technically demonstrate in the context of a conceptual computational model the multicellular behaviour that can emerge when individual cells exhibit basic biologically inspired behaviours such as local cell to cell communication and autoregulation. Biological terms such as cell, molecule, ligand, receptor etc are therefore used to mean conceptual versions of the real thing unless stated otherwise.

### Model and parameters

Each cell can occupy a fixed position in 2D space. Cells are evenly spaced apart and are therefore arranged in a grid like format (fig 2). New cells can be added and old cells removed from any point in space at any point in time, meaning the population size can be changed. However, individual cells cannot move once they are in the population, they are fixed in their location.

**Figure 2 –.**
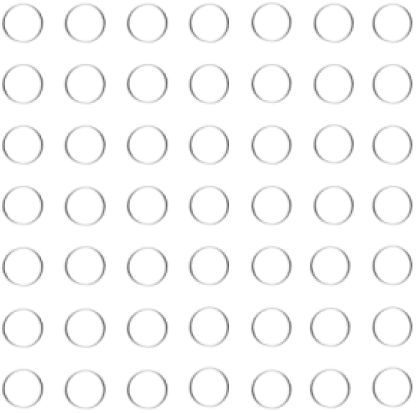
Cells are conceptualised as circles in 2D space. Cells are evenly spaced apart and fixed in their position.

Time exists in discrete timesteps. Cells can send and receive signals at each timestep. A cell can send a signal to any other cell that is directly above, below, to the left or to the right of it, diagonal communication is not possible. A cell cannot send or receive a signal further than its immediate neighbours. When a cell sends a signal, it is propagated in all directions at once (up, down, left, right), this means a cell cannot send different types of signals to different neighbours, only the same type of signal(s) to every neighbour at the exact same time (fig 3).

**Figure 3 –.**
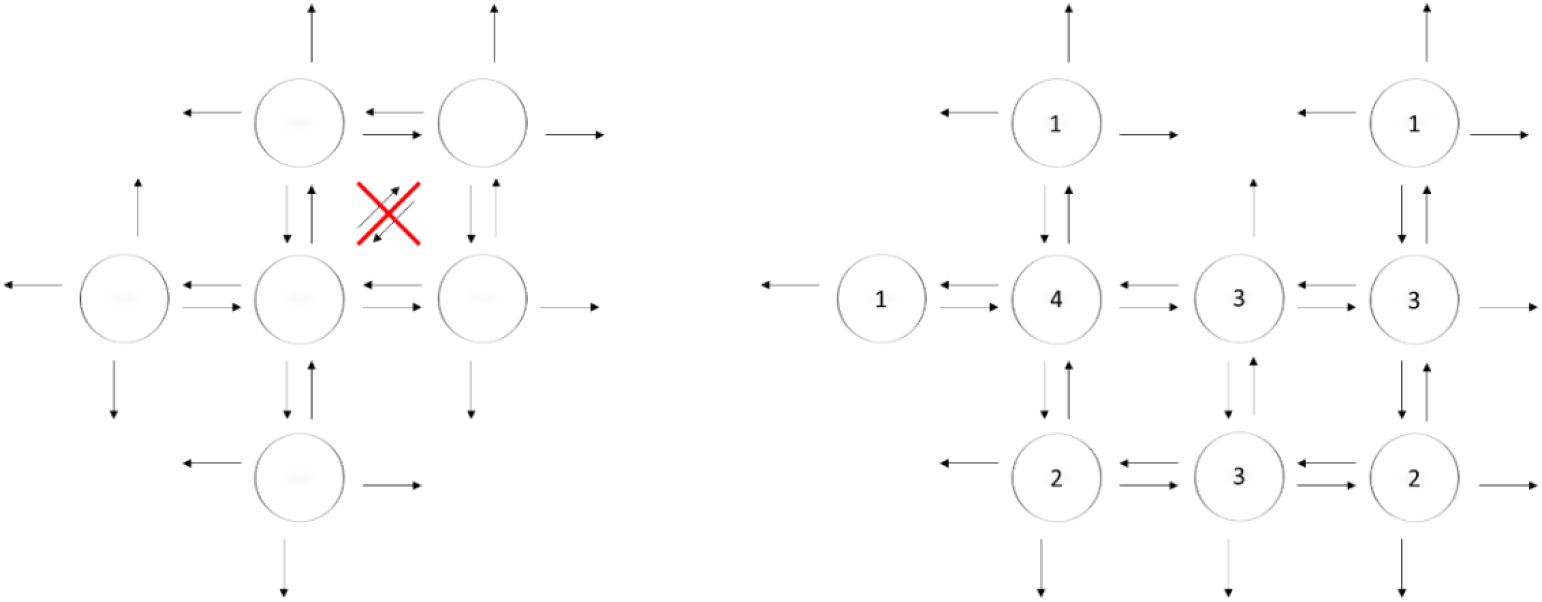
Cells can send signals to any other cell that is directly above, below, to the left or to the right of it, diagonal communication is not possible. A) Arrows represent the signals the cells can and cannot send and receive. B) Schematic of a population of cells sending signals to one another. Numbers inside the cells represent the amount of neighbours each cell has.

It takes one timestep to send and receive signals. In each timestep cells will first receive signals and then send signals to their neighbours. Therefore, signals sent by cells in timestep *x* are received by their neighbours in timestep *x* + 1. Likewise, signals received by cells in timestep *x* are those sent to them by their neighbours in timestep *x* — 1. Signals are transient and act only once upon a cell, i.e. a cell will only sense and process a signal in the timestep it receives it.

In summary, this is an extremely simple hypothetical environment in which every cell is in a fixed grid like position. Cells send signalling molecules at exact rates in all directions at each discrete timestep and these signals are received and processed immediately in the next timestep by their neighbours. Once a signal is received by a cell it is gone, it does not continue to act as a signal to the same or any other cell in subsequent timesteps, ultimately it can be thought of as a one off message/signal sent from one cell to another.

### Cell to cell communication algorithm

This section explains the cell to cell algorithm that enables population size detection and, as a consequence of this, formation of a signalling gradient throughout the population.

Imagine you are a cell in a fixed location, as in the environment constructed above. Your observable world is the 4 cell positions immediately next to you, you cannot receive information from anything beyond these 4 neighbouring cells.

The same applies to all other cells in the population, they have equivalent observable worlds except in a different spatial location.

If all cells emit a single molecule at a constant rate, then what information can a single cell determine about its external environment at any given point in time? It seems it can determine how many neighbours are present, but nothing more. For example if a cell receives 0 molecules in a timestep, it has no neighbours, 1 molecule 1 neighbour, 2 molecules 2 neighbours etc up to the maximum of receiving 4 molecules per timestep meaning it has 4 neighbours and is completely surrounded.

If this applies to every cell, then an individual cell can make no distinction between being surrounded by 4 neighbours when in a small population compared to the same cell being completely surrounded within a larger population (fig 4).

**Figure 4 –.**
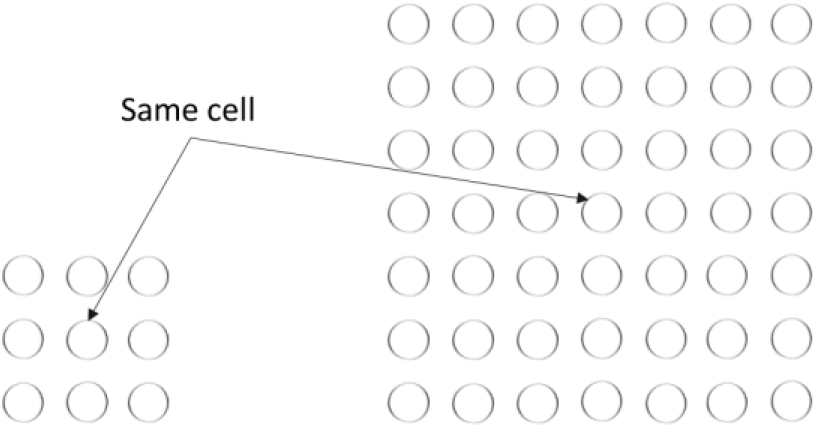
If cells can only send one type of signal at a constant rate then there is no way for individual cells to determine anything about the overall population size. For example, the cell with arrows pointing towards it represents the same cell in different population sizes. The signals it would receive in both populations would be exactly the same if all cells can only send one type of signal at a constant rate, therefore meaning that the highlighted cell would not sense any difference between these 2 population sizes.

How can an individual cell increase its knowledge of the external environment beyond its immediate 4 neighbours? When a cell is completely surrounded by 4 neighbours, it could send out another type of molecule/signal to let its neighbours know that it is surrounded. We can call these different signals ‘mol1’ for the first and ‘mol2’ for the second (fig 5).

**Figure 5 –.**
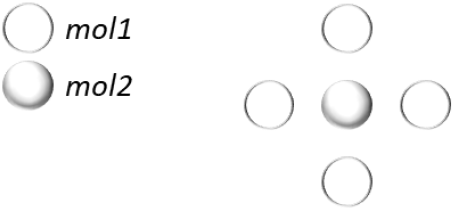
The central cell is surrounded by neighbours and starts sending an additional molecule type named ‘mol2’. Its neighbours are not completely surrounded and therefore send only mol1. The cells are shaded as per the key in the diagram to represent which ‘mol’ type they are sending.

This rule applies to all cells in the population, meaning any cell that is surrounded will send both mol1 and mol2. If any of a cells neighbours are completely surrounded by their own neighbours then it will receive mol2 from that neighbour (fig 6).

**Figure 6 –.**
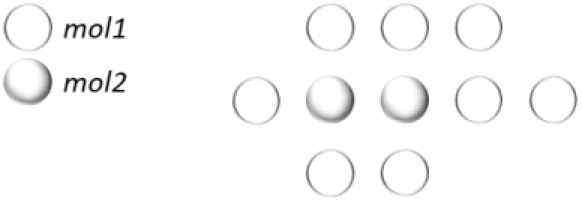
Two cells in this population are completely surrounded by neighbours and therefore sending mol1 and mol2. Cells that are not surrounded continue sending only mol1.

If all of the neighbours of a surrounded cell became surrounded themselves, and therefore began sending mol2, the centre cell would receive the maximum amount of mol2 possible per timestep, which is 4, one from each of its 4 neighbours. From this the central cell can deduce that its neighbours are also surrounded based on the fact that they are all sending it mol2, and it can therefore deduce information about its external environment beyond its limited observable world of its immediate 4 neighbours (fig 7).

**Figure 7 –.**
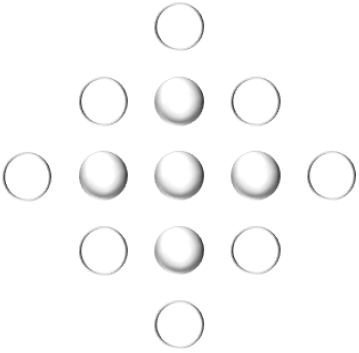
The centre cell is surrounded by neighbours which are also surrounded by neighbours. The centre cell can deduce that all its neighbours are surrounded from the fact that it is receiving mol2 from all of them.

This basic signalling rule can be reapplied each time all of a cell’s neighbours are producing the same signal as itself, enabling the centre most cell in a population to accurately determine the overall population size. For example, once the centre cell is receiving 4 mol2 per timestep (meaning all of its neighbours are surrounded by neighbours themselves and therefore sending mol2), it can send a third type of molecule/signal which can be called mol3 (fig 8). If all of the centre cells neighbours then became surrounded by neighbours that themselves are surrounded by neighbours (in other words, the neighbours of neighbours of the centre cell are also surrounded by neighbours), then the centre cells immediate neighbours will also send mol3 (because they will be receiving 4 mol2 per timestep) and the centre cell will receive 4 mol3 per timestep, enabling it to deduce that its neighbours are surrounded by neighbours that are again surrounded by neighbours. We can continue this process indefinitely by adding new ‘mol’ types so that the centre cell can always deduce the population size, no matter how large (fig 9).

**Figure 8 –.**
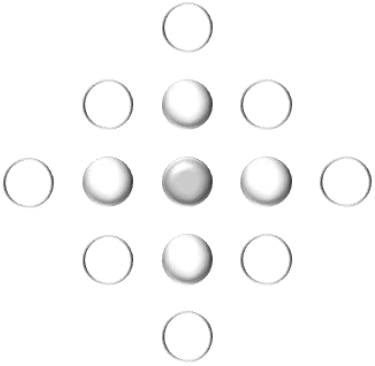
The centre cell is surrounded by neighbours which are also surrounded by neighbours. This means the centre cell will now start sending an additional molecule type named ‘mol3’.

**Figure 9 –.**
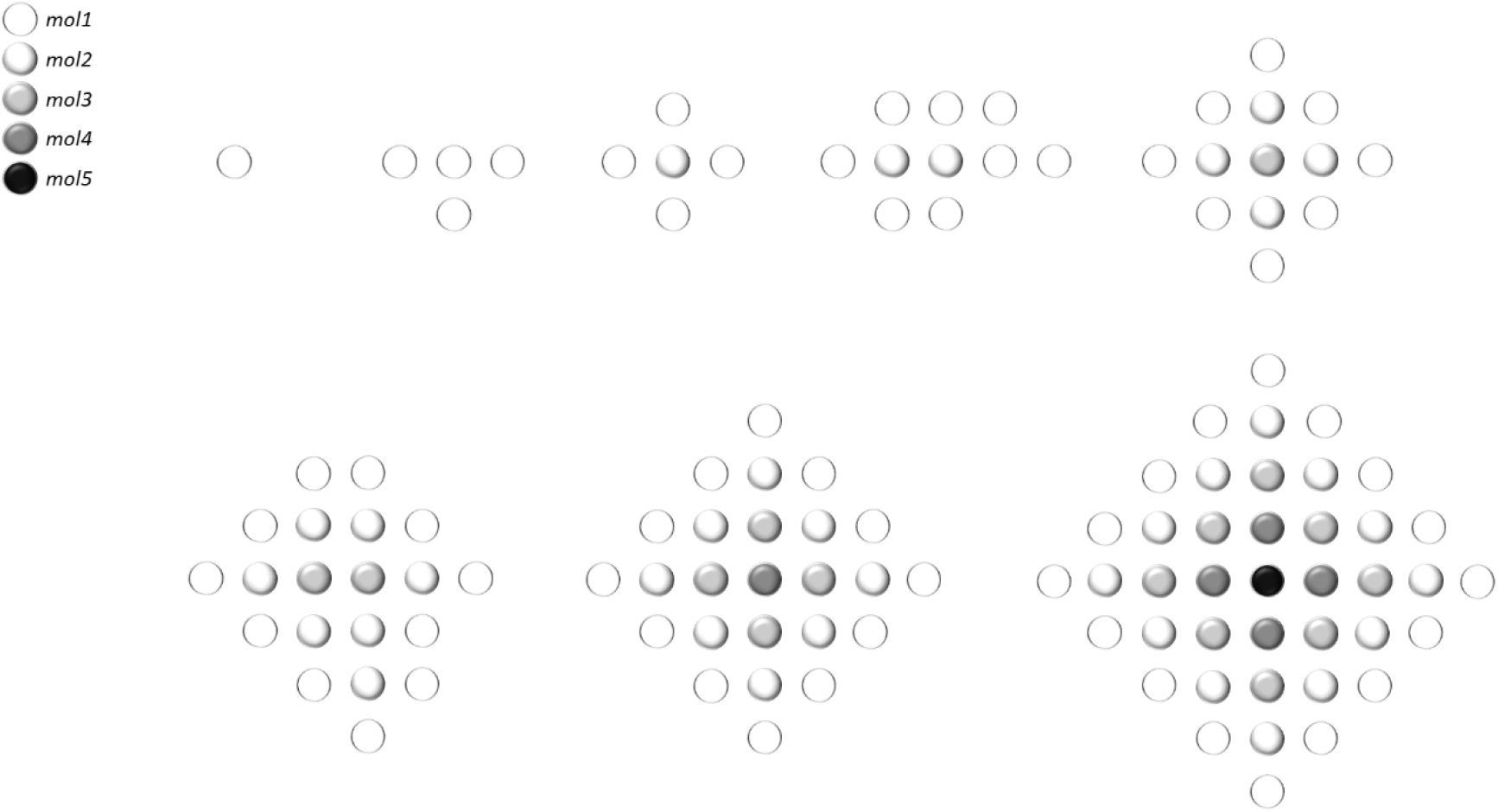
Stable signalling patterns at various population sizes. When a cell is surrounded by neighbours that are all sending mol*x* it will send mol*x* + 1.

The communication algorithm is straightforward: when all of your neighbours are sending mol*x* then send mol*x* + 1.

As a consequence of this, a ‘gradient’ of molecule types (mol1, mol2, mol3, etc) emerges in the overall cell population, with the highest number mol type in the centre and lowest at the periphery.

### Gradient generation with one molecule type

It is commonly observed in real biological systems that a gradient of a single molecule type spreads across a population of cells (Blair 2007, Hill 2017, O’Connor et al 2006, Placzek & Briscoe 2018). Here different ‘mol’ types are being used as opposed to a single molecule. However, if the change in each molecule type is translated to an increase in the rate of production of the same molecule, then the same or similar results can be generated with one molecule, which is that there will be a gradient of rates of molecular signalling/production across the population, with high rates in the centre and low rates at the periphery.

The rule used above can be modified so that equivalent behaviour occurs but using only one molecule. For example, instead of saying:

> *“When a cell is surrounded by neighbours that are all sending molx then the surrounded cell sends molx* + 1”.

Equivalent behaviour with one molecule can be achieved by saying:

> *“When a cell is surrounded by neighbours that are all sending greater than or equal to x molecules per timestep then the surrounded cell will send x + 1 molecules per timestep”*.

If for each cell *x* = the minimum value being produced out of all its neighbours, then the above statement will behave in an exactly equivalent manner to the preceding statement. The Implementation section of this document shows examples of gradient formation using this algorithm.

### Formal definition

This section defines the fundamental properties of the model that is used in all examples in this paper, which is based on the conceptual model described in the previous section. Any other properties of the model that are specific to a particular example are defined in the relevant section.

The model is defined for cells organised in a 2-dimensional grid as in the example in the previous section but also for cells in a 1-dimensional row. It is simpler to demonstrate and visualise the process of gradient formation with 1D because each timestep can be viewed as a row and directly compared to the previous row/timestep.

To clarify, from here on in when referring to the *‘amount of molecules produced by a cell in a timestep’* or similar terminology, this is taken to mean the amount of molecules a cell sends in each direction in a particular timestep, including to itself. The absolute value produced will therefore be this value multiplied by the maximum amount of neighbours a cell can have plus one. For example, if a cell has a maximum of 4 neighbours as in the previous section, then if we say a cell produces *x* molecules in a timestep, this value *x* is sent to all the cell’s neighbours plus itself, so the absolute or total value produced by the cell would actually be 5*x*.

#### 2-Dimensional

There are 2 matrices *C* and *G* of size *trc*. Where the t dimension represents each timestep and *r* and *c* represent the *x* and *y* axis of 2D space respectively.

*C* is a binary matrix that represents the physical location of cells in 2D space. Each *C_zxy_* = 1 represents a cell and each *C_zxy_* = 0 represents empty space.

The neighbours of cell *C_zxy_* would therefore be *C*_*z,x*−1,*y*_, *C*_*z,x,y*+1_, *C*_*z,x*+1,*y*_, *C*_*z,x,y*−1_

*G* represents the signals produced by cells. Signals sent by cell *C_zxy_ = G_zxy_*, which means the signals cell *C_zxy_* will receive in timestep *z* are those produced by itself and its neighbours in the previous timestep *z* − 1:

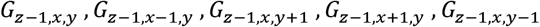

The first term above *G*_*z*−1*x,y*_ represents a cell receiving its own signal, meaning that a cell will self-regulate and continue to produce molecules in the absence of any neighbours. The other 4 terms represent the signals received from each neighbour.

Signals produced by cells are a function of the signals received in the same timestep, meaning each *G_zxy_* is defined as:

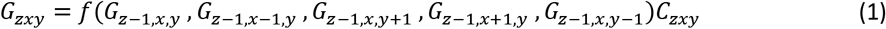

The function is multiplied by the binary value in *C_zxy_* so that a positive non-zero value can be produced only if a cell exists in that physical location in space, which is represented by *C_zxy_* = 1, and empty space is represented by *C_zxy_* = 0.

The function *f* is defined separately for each specific implementation in this paper.

#### 1-Dimensional

For 1D, the core difference is that the *y* axis in the 2D space is lost, so cells and signals now exist only along the *x* axis and are represented by the *r* dimension in matrices *C* and *G* respectively. These are effectively vectors each of which corresponds to a timestep *z* in the *t* dimension (fig 10).

**Figure 10 –.**
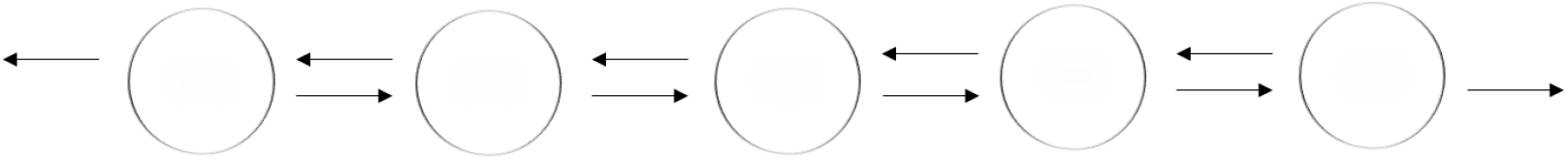
The 1D model is the same as the 2D model except that cells are arranged in a row instead of 2D grid. They are evenly spaced apart and fixed in their position. Cells can send signals to any other cell that is directly to the left or to the right of it.

There are 2 matrices *C* and *G* of size *tr*. Where the *t* dimension represents each timestep and *r* represents the 1-dimensional row of cells at each timestep.

*C* is a binary matrix that represents the physical location of cells in a 1D row. Each *C_zx_* = 1 represents a cell and each *C_zx_* = 0 represents empty space.

The neighbours of cell *C_zx_* would therefore be *C*_*z,x*−1_, *C*_*z,x*+1_

*G* represents the signals produced by cells. Signals sent by cell *C_zx_ = G_zx_* and cell *C_zx_* will receive signals from:

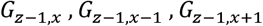

The first term above *G*_*z*−1,*x*_ represents a cell receiving its own signal, meaning that a cell will self-regulate and continue to produce molecules in the absence of any neighbours. The other 2 terms represent the signals received from each neighbour.

Signals produced by cells are a function of the signals received in the same timestep, meaning each *G_zx_* is defined as:

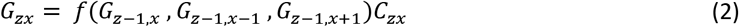

The function is multiplied by the binary value in *C_zx_* so that a value will be produced only if a cell exists in that physical location in space, which is represented by *C_zx_* = 1, and empty space is represented by *C_zx_* = 0.

The function *f* is defined separately for each specific implementation in this paper.

#### Initial conditions

In all implementations in this paper including those in the appendix, timestep 1 is taken as the initial conditions and therefore the production values of cells are set manually. All subsequent timesteps use the equations defined in the model to determine the production rate of each individual cell.

Some cells are initially set to produce zero molecules so that they will be dormant until a signal is received by one of its neighbours. Equation (1) and (2) on their own will result in cells always producing at least some base amount of molecule even if they are initially set to zero because self-regulation is incorporated into the equation. Whilst this is not a problem, it is interesting to see gradients form from one or a subset of initially producing cells and then have this propagate out into other cells which start signalling only when signals reach them. Therefore, a condition is incorporated into the model which is that cells will only perform the autoregulation function and start producing when 1 or more of their neighbours are producing. The condition is defined as:

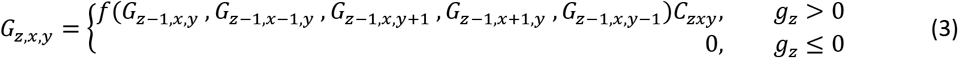

For 2D and

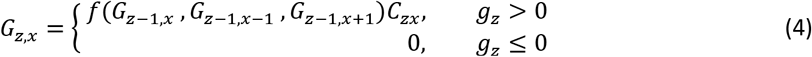

For 1D.

Where *g_z_* equals the sum of all molecules produced by a cell and its neighbours in the previous timestep, meaning:

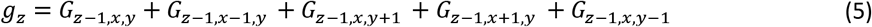

For 2D and

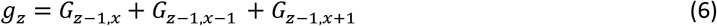

For 1D.

### Implementation

The model has been implemented using Microsoft Excel, whereby the equations can be entered as formulas in each cell and the formation of the gradient through time can be viewed as static rows or grids of numbers for 1D and 2D respectively (figure 11).

**Figure 11 –.**
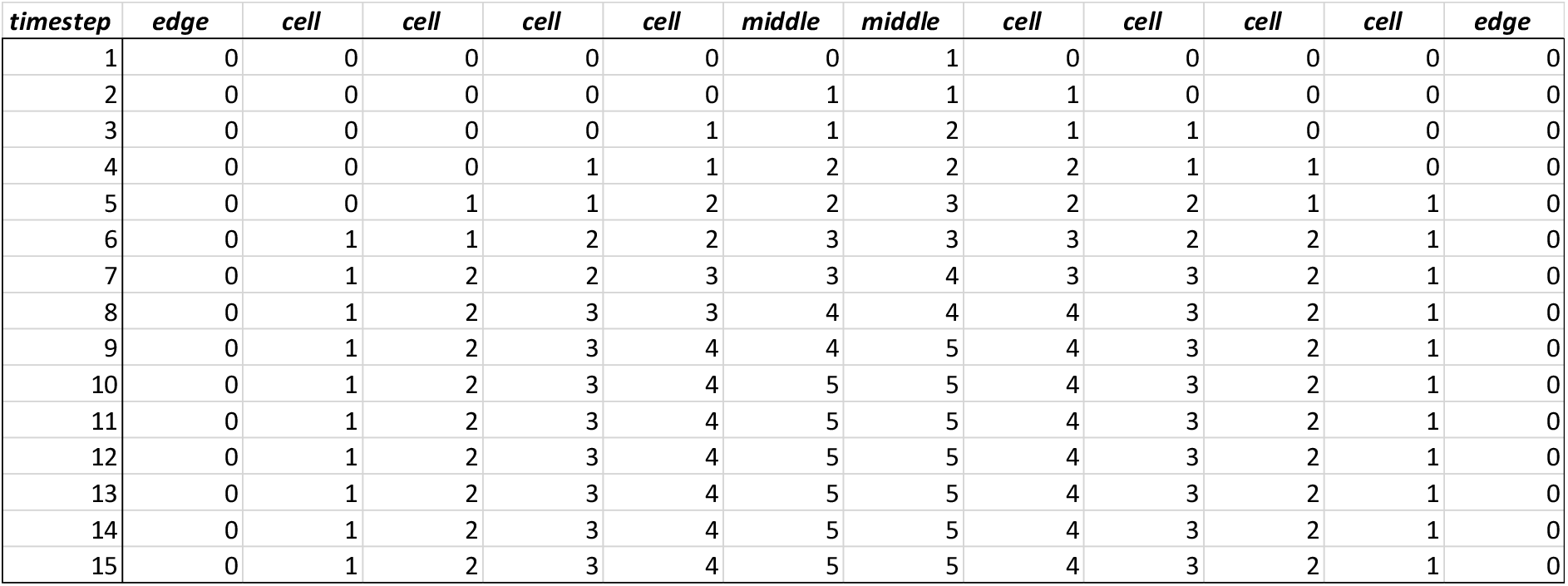
Table showing the formation of a gradient over time using the autoregulation rule defined in equation (*8*) where *a* = 1. Each row represents a timestep and the columns represent cells. The edge columns are not cells but represent empty space. The number under each cell column represents the amount of molecules that cell sends to each of its neighbours and itself in that timestep. A cell receives molecules produced by each of its neighbours and itself in the previous timestep. In accordance with the rule defined in equation (*8*), at each timestep cells produce the minimum value of that produced by their neighbours in the previous timestep plus 1. The gradient stabilises at timestep 10, meaning that in all subsequent timesteps cells will continue to produce at the same rate.

#### Autoregulation Function

To mimic the behaviour described in the signalling concept section, the function *f* in equation (1) must implement the following statement:

> *“When a cell is surrounded by neighbours that are all sending greater than or equal to x molecules per timestep then the surrounded cell will send x + a molecules per timestep”*.

Which can be done as follows:

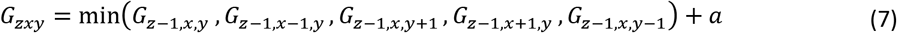

For 2D and

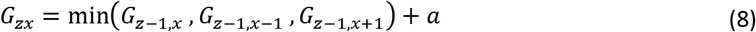

For 1D. Where *a* is a real number. The min () function takes the minimum value of all those listed in the brackets.

This function means a cell will simply take the value received from its lowest producing neighbour (including neighbours producing zero or empty space) and add *a*. All the examples below are done using *a* = 1. This means *x* in the statement above is always equal to the value of molecules a cell receives from its lowest producing neighbour.

#### Examples

Figure 11 shows the formation of a gradient in a row of 10 cells.

Figure 12 shows the formation of a gradient in a row of 30 cells and the gradient that is formed in a 10 by 10 grid and 30 by 30 grid of cells. To save space only the starting condition and stable state are shown for 2D models.

**Figure 12 –.**
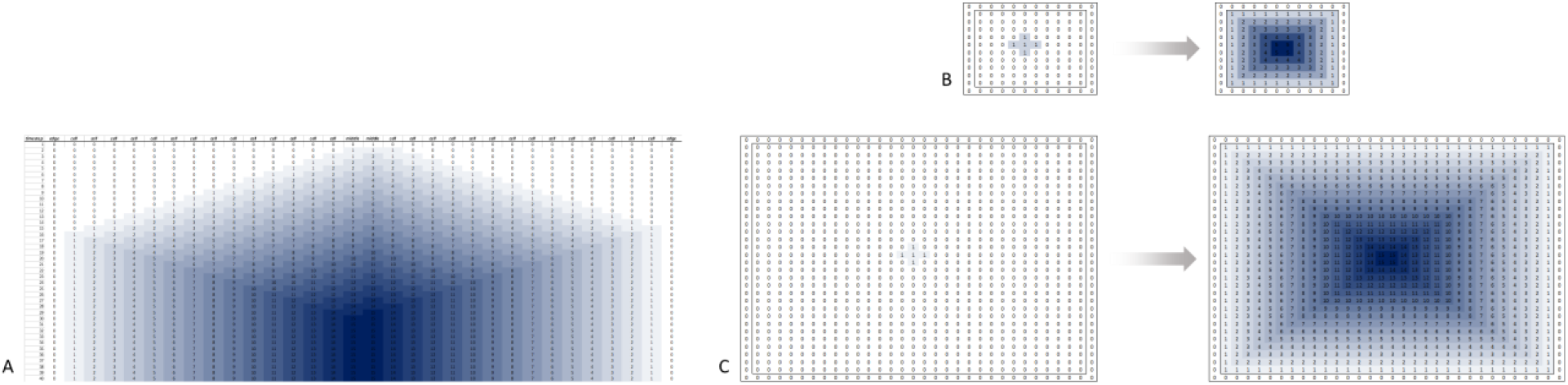
A) table showing the formation of a gradient over time in a row of 30 cells using the autoregulation rule defined in equation (*8*) where *a* = 1. Columns are shaded different strengths of blue to indicate the relative production rate of cells. The gradient stabilises at timestep 30. B) 10 by 10 grid of cells showing the formation of a gradient using the 2D model as defined in equation (*7*) where *a* = 1. Only the initial condition and stable state are shown. The gradient stabilises at timestep 11. C) 30 by 30 grid of cells showing the formation of a gradient using the 2D model as defined in equation (*7*) where *a* = 1. Only the initial condition and stable state are shown. The gradient stabilises at timestep 31.

Figure 13 shows a gradient forming in a row of 50 cells, with various perturbations made throughout the course of development. The perturbations are uneven initial conditions, overproducing cells and population size changes. The stability of the algorithm is evident from this example, as with each perturbation the gradient always restabilises.

**Figure 13 –.**
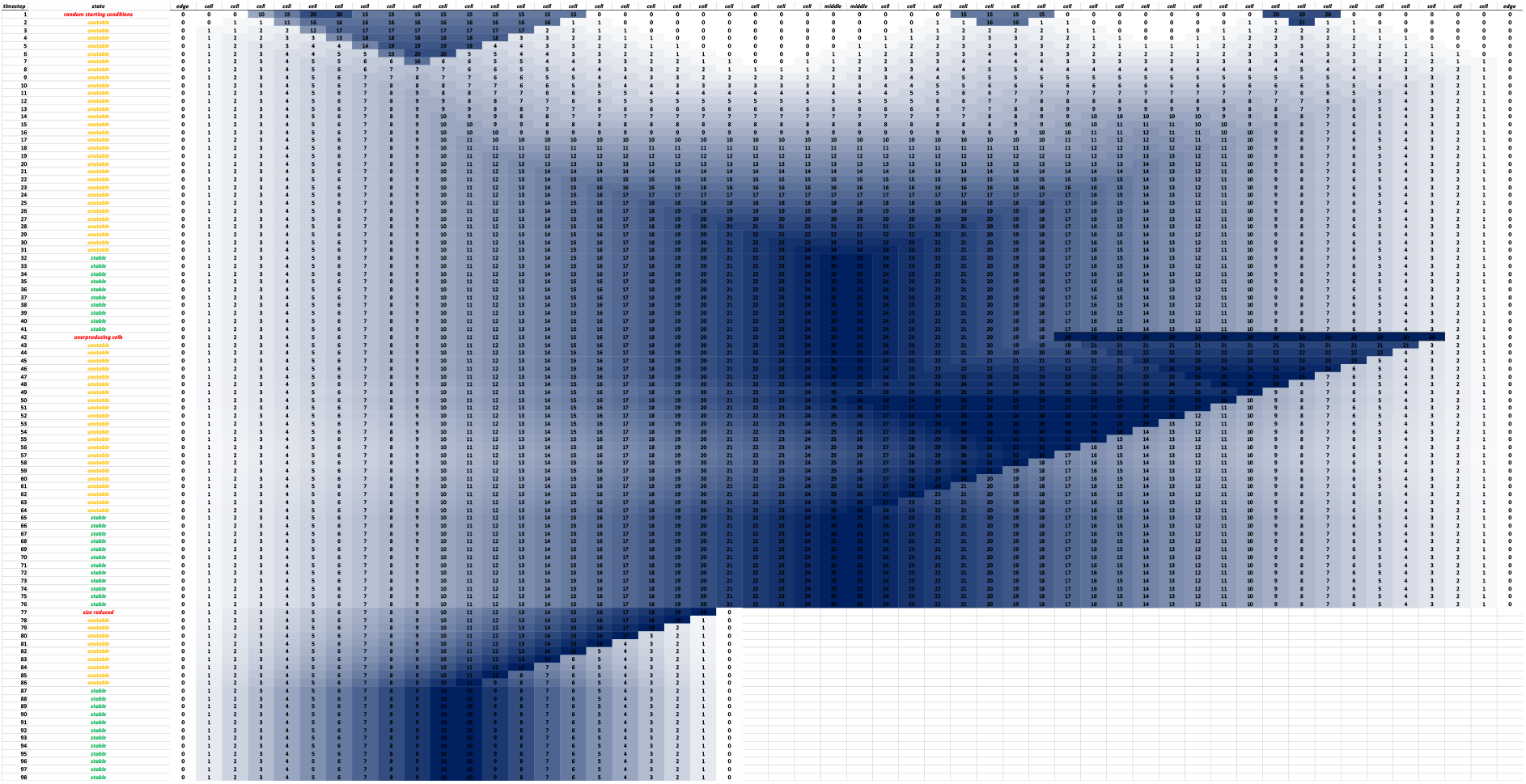
Table showing the formation of a gradient over time in a row of 50 cells using the autoregulation rule defined in equation (*8*) where *a* = 1. Cells are shaded different strengths of blue to indicate the production rate. The blue colouring of cells is used to help visualisation of the gradient formation, therefore the strength of blue indicates the relative signalling rate of cells within the population at each timestep, rather than the absolute value. For example, in the smaller gradient the central cells are producing a lower absolute value of signal than in the larger gradient but the strength of blue is the same. The state column describes the state or activity of the population: ‘Unstable’ means the gradient is changing and moving towards a stable state. ‘Stable’ means the signalling rates of all cells is stable, such that in all subsequent timesteps every cell will produce the same amount of signalling molecules unless the population is perturbed in some way. Other content in the State column are various perturbations that occur on that timestep, these are Random starting conditions, Over producing cells and Size reduction. It can be seen that after each perturbation the gradient eventually restabilises.

Figure 14 shows a gradient forming in a 30 by 30 grid of cells with uneven initial conditions. The stability of the algorithm for 2 dimensions is the same as that for 1 dimension, it is robust to perturbations and will always stabilise back to a consistent gradient.

**Figure 14 –.**
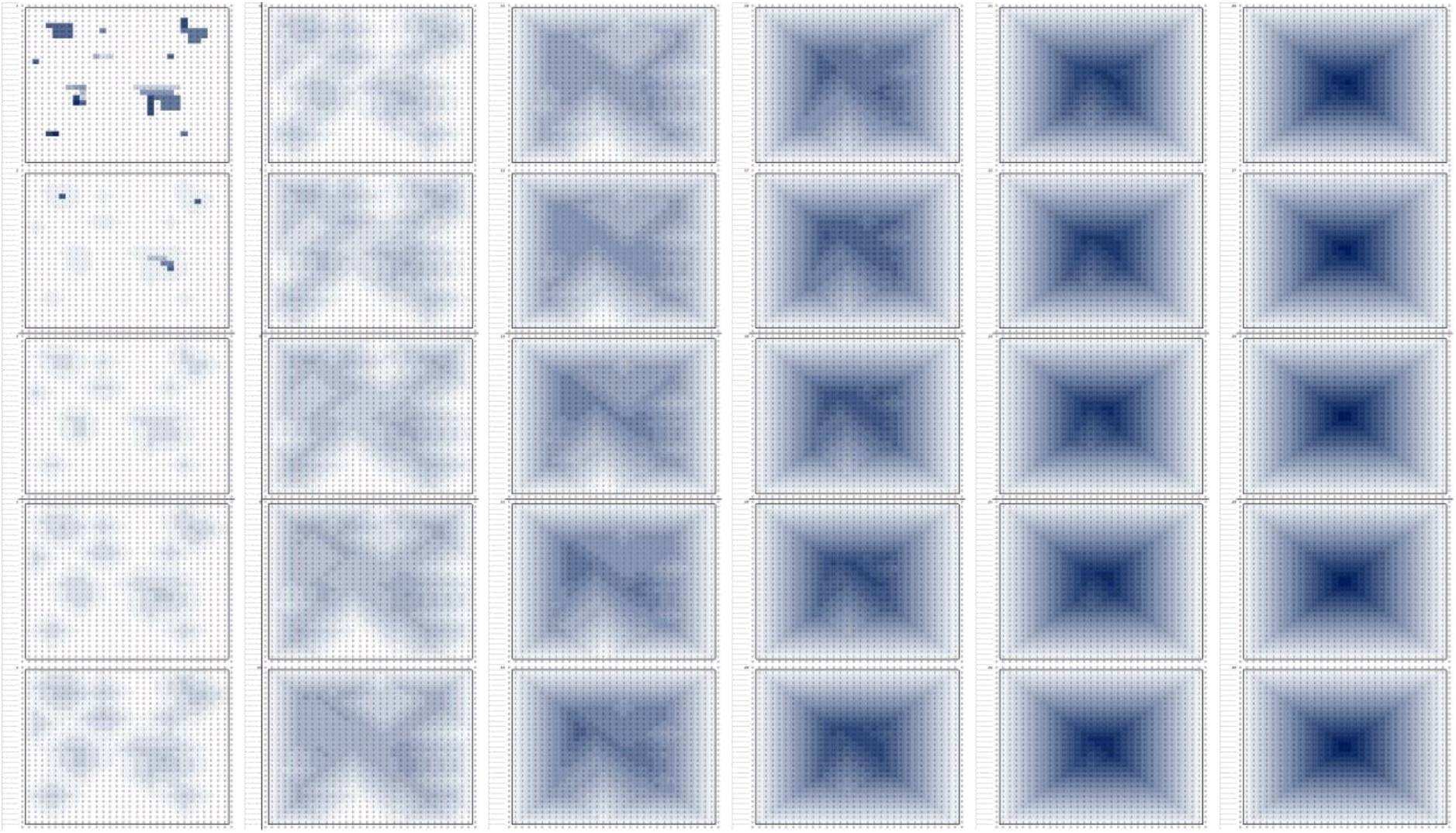
Formation of a gradient over time with uneven initial conditions in a 30 by 30 grid of cells using the autoregulation rule defined in equation (*7*), where *a* = 1. The strength of blue indicates the rate of signalling for each cell, the darker the blue, the faster the rate of signalling of that cell. Moving from top to bottom then left to right, each image shows the production rates of cells at each timestep. 30 timesteps are shown. The gradient stabilises at timestep 28, meaning cells exhibit the exact same signalling behaviour in subsequent timesteps.

In both the 1D and 2D examples with perturbations, overproducing cells are downregulated very quickly, this is because their neighbours are producing less than them and the rule defined in equation (8) states that a cell will always produce the lowest value received out of itself and all its neighbours plus 1.

### Discussion and relevance to biology

There are extremely obvious differences between the model used in this paper and real biological systems, of course real cells do not maintain exact positions in space, equidistant from one another and signal at perfectly constant rates. What is arguably still interesting about the results is the ease with which a gradient can be formed and maintained using very simple rules, especially given that one of the fundamental signalling behaviours required to achieve this is one which is prevalent in real biological systems, autoregulation. The other core aspect of the model is that the rate at which cells regulate their signalling occurs in accordance with the signalling rates of their neighbours.

The gradient formation mechanism shown is highly robust to perturbations, however it is not robust to noise. The robustness to perturbations is reliant on cells always signalling correctly. Real biological systems are far more stochastic and are therefore likely to incorporate signalling rules that tolerate a high level of noise. It is conceivable that some similar form of the algorithm used in this model would be more noise tolerant by using larger values for certain parameters, such as signalling rates, which could then allow less precise thresholds required to induce autoregulatory signalling changes and enable tolerances for variations or perturbations in signalling.

The signalling algorithm used to generate gradients, as defined in equation (7) and (8), requires cells to distinguish between signalling rates of each of their neighbours. It would be surprising if real biological cells are able to do this in the context of responding to morphogens. This is because the signalling mechanisms typically involve secretion of signalling molecules into the extracellular environment (Rogers and Schier 2011, Sagner and Briscoe 2017) where they can freely diffuse, from this it does not seem feasible that a cell could determine which of its neighbours produced any particular signalling molecule that it receives. In addition to this, at any point in time cells may be moving, differentiating, undergoing apoptosis or other activities which mean individual cells are unlikely to always have the same immediate neighbours. Even though this is the case, it still seems plausible that the molecular mechanisms cells use to regulate their signalling could be such that they behave in a way which indirectly causes them to signal in accordance with the signalling behaviour of their neighbours, without having to actually know the exact behaviour of their neighbours. Appendix 1 demonstrates how this can be achieved using a combination of biologically inspired regulatory mechanisms such as ligand and receptor complexes, ligand inhibitors and receptor regulation. This allows cells to simply respond to absolute quantities of signalling molecule they receive without regard for the exact behaviour of their neighbours, yet still behave in a manner which is concurrent with signalling behaviour of their neighbours and similar to that defined in equation (7) and (8). This is only intended to demonstrate that this is feasible in the context of the model used in this paper, it is not making any claim that real biological systems work in this way.

Appendix 2 shows a method of implementing opposing gradients that scale according to overall population size. This is inspired by the fact that morphogen gradients in real biological systems can scale according to population size (Almuedo-Castillo et al 2018, Collins et al 2018, Inomata 2017), once again however this has only been done to demonstrate feasibility of scale invariance in the context of the model used in this paper and without any claim that real biological systems adopt the same or similar mechanisms. The scale invariance section in Appendix 2 involves a discussion about the information that individual cells require in order to achieve scale invariance. It also discusses the issue of signalling synchronicity, whereby signalling alterations in one part of a population must have sufficient time to propagate throughout the rest of the population so that cells make decisions based on signals that are providing accurate information.

## Appendix 1 – Gradient stabilisation using mechanisms inspired by biology

This section shows how biologically inspired regulatory mechanisms such as ligand and receptor complexes, ligand inhibitors and receptor production and degradation can be used to generate gradients in a similar manner to that using the function defined in equation (7) and (8).

This section is only intended to show ways in which biologically inspired mechanisms can be used to achieve certain types of regulatory behaviour in the context of the conceptual model being used in this paper. It is not making any claim that real biological systems work in this way.

The different regulatory methods generate gradients that vary in terms of stability. It begins with the least stable and ends with the most stable. The final method incorporates all the preceding regulatory mechanisms to achieve greater robustness.

### Upregulation

This is a highly simplistic rule in which cells upregulate the absolute value of molecules received from their neighbours and itself:

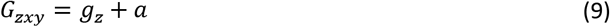

Where *a* is a real number. Recall that *g_z_* equals the sum of all molecules produced by a cell and its neighbours in the previous timestep as defined in equations (5) and (6).

This means that the quantity of molecules produced by a cell in timestep *t* is equal to the sum of all molecules received in timestep *t* plus an additional amount *a*.

It is also possible to have a multiplicative version:

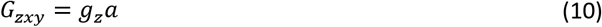

This means that the quantity of molecules produced by a cell in timestep *t* is equal to the sum of all molecules received in timestep *t* multiplied by a factor *a* (fig 15).

**Figure 15 –.**
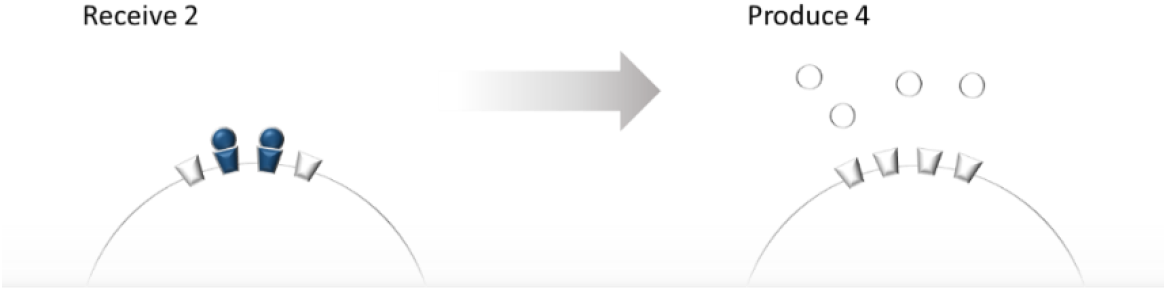
Conceptual diagram of upregulation using equation (*10*) where *a* = 2. The cell receives 2 molecules and subsequently produces 4, which is the amount received multiplied by 2.

In both the additive and multiplicative versions, the production rate of each cell continues to increase at each timestep and will continue towards infinity at each successive timestep. This is not desirable because the gradient will never stabilise and cells will be unable to determine the population size. In order to determine population size, the signalling behaviour of all cells in the population needs to reach a certain stable rate of production that corresponds to a particular population size.

Figure 16 shows a step by step generation of a gradient in a 1D row of cells using the additive upregulation rule.

**Figure 16 –.**
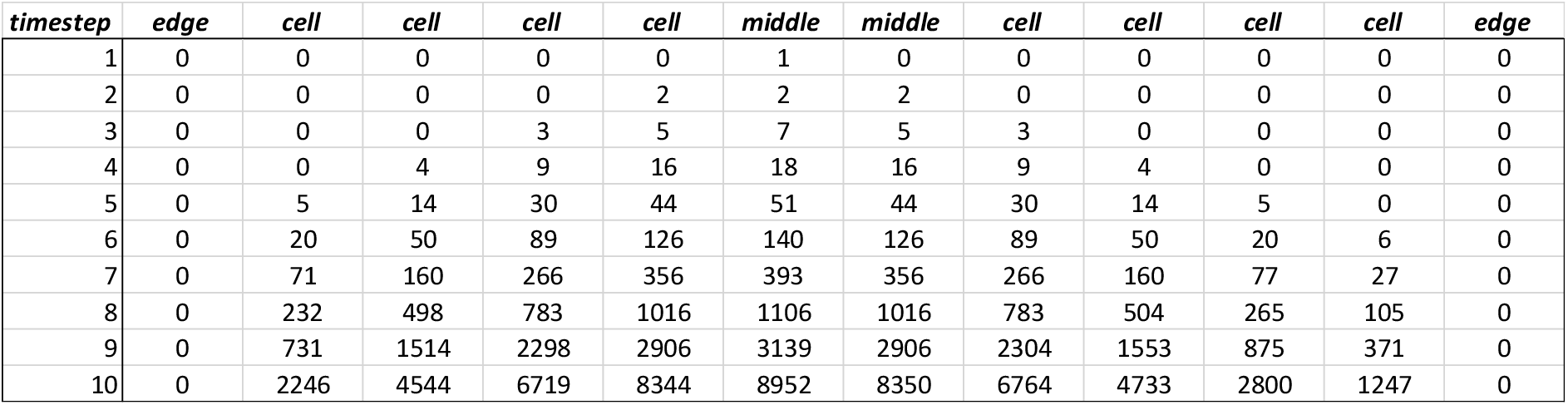
Table showing the formation of a gradient over time using the additive autoregulation rule defined in equation (*9*) where *a* = 1, meaning a cell produces the amount of molecules it receives at each timestep plus 1. Each row represents a timestep and the columns represent cells. The edge columns are not cells but represent empty space. The number under each cell column represents the amount of molecules that cell sends to each of its neighbours in that timestep. A cell receives the sum of molecules produced by each of its neighbours and itself in the previous timestep.

Figure 17 shows a step by step generation of a gradient in a 1D row of cells using the multiplicative autoregulation rule.

**Figure 17 –.**
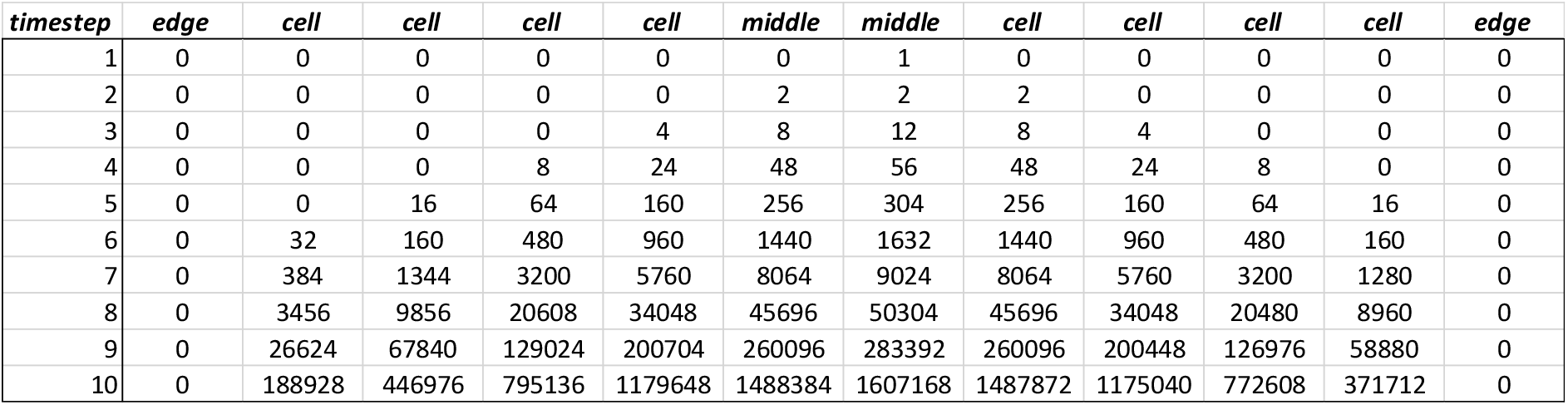
Table showing the formation of a gradient over time using the multiplicative autoregulation rule defined in equation (*10*) where *a* = 2, meaning a cell produces twice the amount of molecules it receives at each timestep. Each row represents a timestep and the columns represent cells. The edge columns are not cells but represent empty space. The number under each cell column represents the amount of molecules that cell sends to each of its neighbours in that timestep. A cell receives the sum of molecules produced by each of its neighbours and itself in the previous timestep.

Figure 14 shows a conceptual view of cell signalling behaviour when the multiplicative autoregulation rule is used and *a* = 2.

**Figure 18 –.**
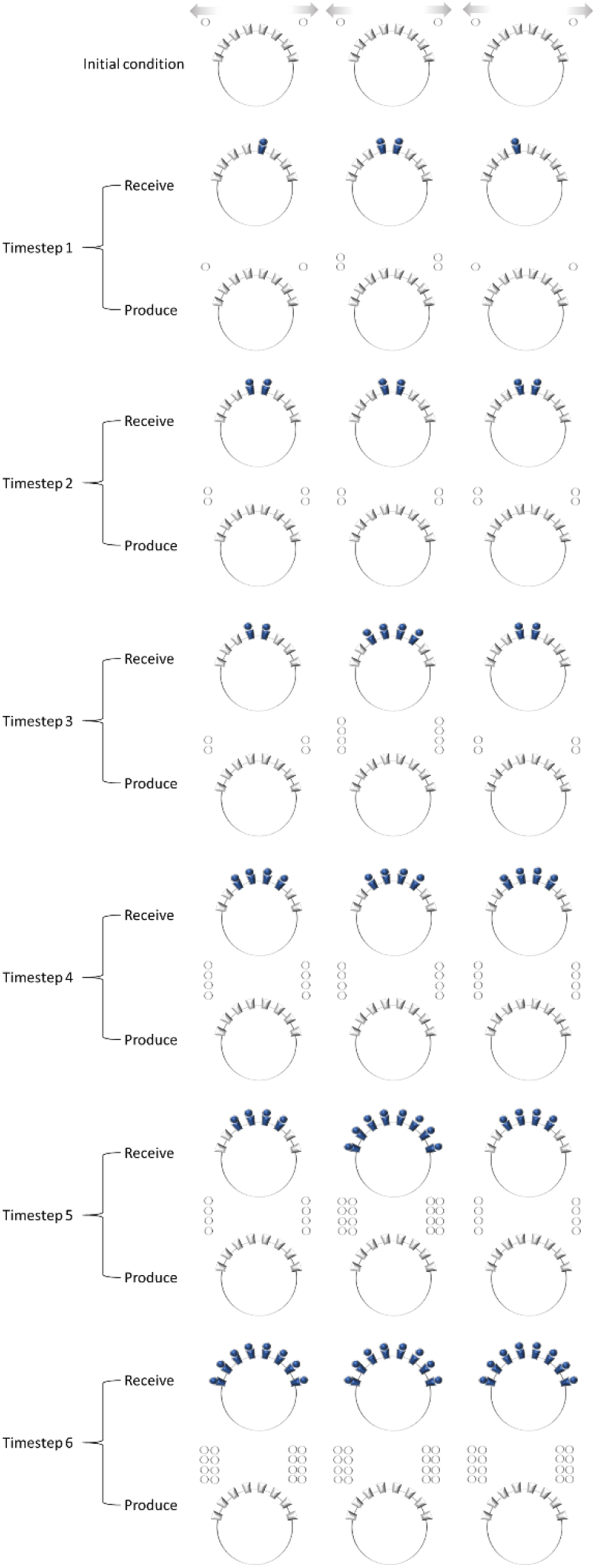
Conceptual diagram showing the signalling behaviour of cells that are using the multiplicative autoregulation rule defined in equation (*10*) where *a* = 2. Each timestep involves cells receiving then producing molecules. In the receive steps, receptors are coloured blue when a signalling molecule binds to them and this represents a cell receiving a molecule. In the produce steps, the molecules shown on either side of each cell are what is sent to neighbouring cells on the same side, and will be received by their neighbours in the next timestep. Self-regulatory signals that cells send to themselves are ignored in this diagram, which does not actually affect the fundamental properties of the multicellular behaviour.

### Molecular complexes

Stabilisation can be achieved if there is a threshold value of molecules that must be received in a single timestep in order for a cell to produce at a certain rate. The threshold value should ideally correspond to the amount of molecules a cell will receive when it is surrounded by neighbours that are all producing at a rate greater than or equal to itself.

The conceptual example used in the main text is in effect using this type of threshold mechanism. Once a cell is surrounded it will receive the threshold amount of molecules needed to produce the next ‘mol’ type. It will not produce the next ‘mol’ type until the threshold value is received. Any value of molecules received above one threshold but still below the next will not cause any change in signalling. The cell is only responding to step changes in value of molecule received.

A similar threshold type mechanism using a single molecule can be created if cells only respond to integer divisions of the overall amount of molecules received. This is done by saying that a cell produces *x* molecules for every *n* received. For example if a cell produces 1 molecule for every 3 received, then if a cell received say 6 molecules it will produce 2, and if it received 7 or 8 molecules it would still produce 2. Only once it received 9 molecules would it increase to produce 3. A molecular mechanism for achieving this that is biologically inspired is that molecules must form complexes of certain integer values in order to bind to and activate a receptor (fig 19).

**Figure 19 –.**
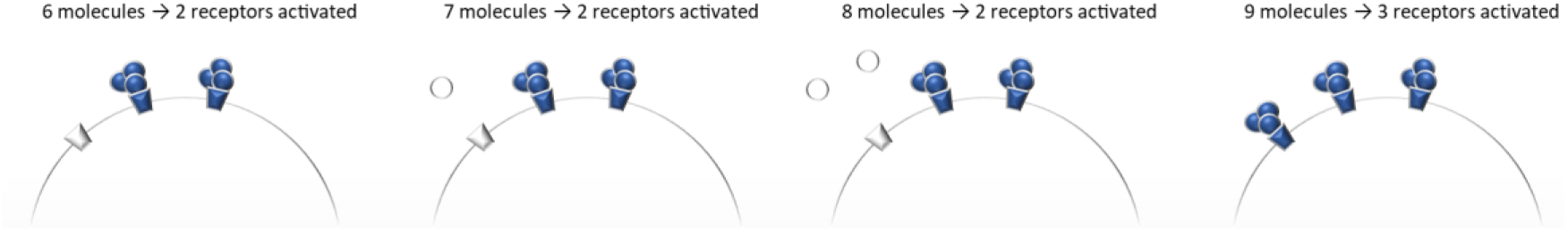
Conceptual diagram demonstrating how threshold values of molecules are required in order to activate certain amounts of receptors if molecules must form a trimer in order to bind to a receptor. Moving from left to right, the cell receives 6, 7, 8 and 9 molecules respectively. Only when the cell receives 9 molecules are 3 receptors activated, in all other cases only 2 receptors are activated.

The following function can be used to implement this autoregulation rule. The floor function ⌊ ⌋ rounds numbers down to the nearest integer:

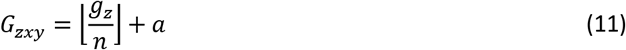

If *n* = maximum number of neighbours a cell can have plus one (the plus one accounting for molecules received from itself), then this results in a cell needing to be surrounded by neighbours that are producing at the same rate as itself in order for its own rate of production to increase. The division by *n* and floor function corresponds to saying produce *x* for every *y* received. The + *a* term is the amount by which a cell will increase its production rate when it is surrounded by neighbours that are producing at the same rate as itself.

Figure 20 shows a step by step generation of a gradient in a 1D row of cells using the autoregulation rule defined in equation (11).

**Figure 20 –.**
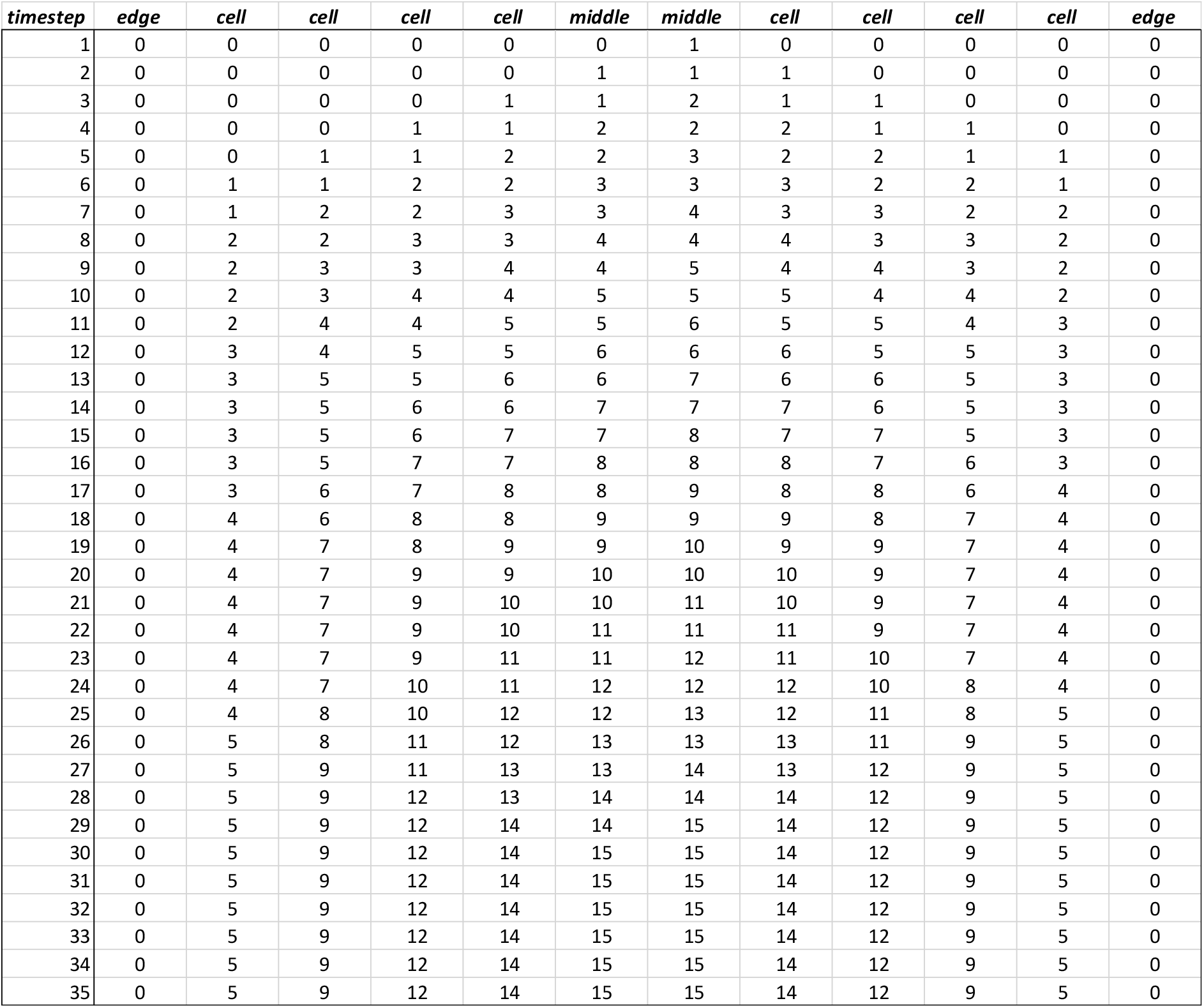
Table showing the formation of a gradient over time in a row of 10 cells using the autoregulation defined in equation (*11*), where *a* = 1 and *n* = 3, meaning a cell produces 1 molecule for every 3 received plus 1. Each row represents a timestep and the columns represent cells. The edge columns are not cells but represent empty space. The number under each cell column represents the amount of molecules that cell sends to each of its neighbours in that timestep. A cell receives the sum of molecules produced by each of its neighbours and itself in the previous timestep. The gradient stabilises at timestep 30, meaning that in all subsequent timesteps cells will continue to produce at the same rate.

Figure 21 shows equivalent behaviour to figure 20 but for larger population sizes and 2D populations.

**Figure 21 –.**
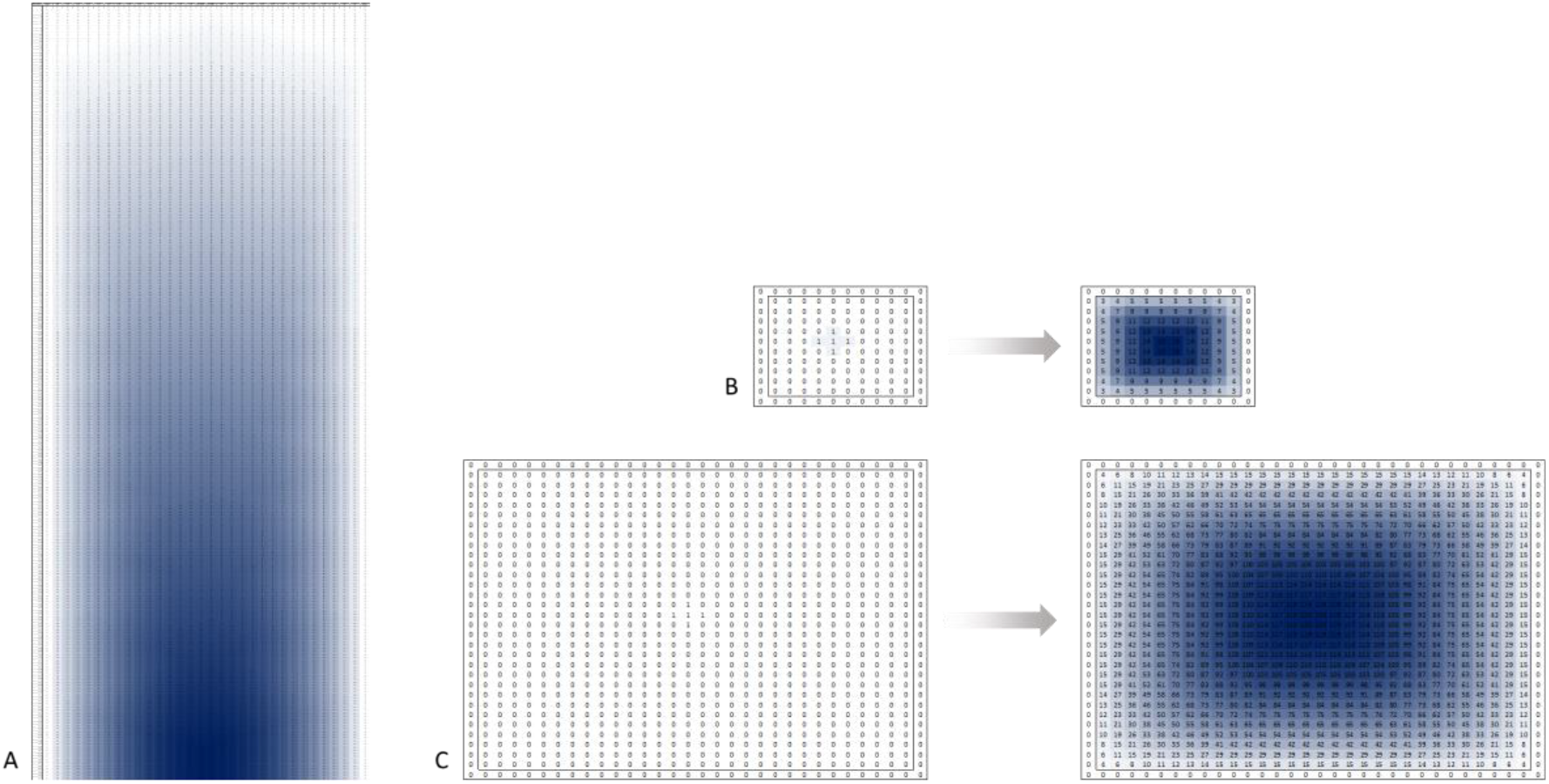
A) table showing the formation of a gradient over time in a row of 30 cells using the autoregulation rule defined in equation (*11*), where *a* = 1 and *n* = 3, meaning a cell produces 1 molecule for every 3 received plus 1. Cells are shaded different strengths of blue to indicate the production rate. The gradient stabilises at timestep 240. B) 10 by 10 grid of cells showing the formation of a gradient using the autoregulation rule defined in equation (*11*), where *a* = 1 and *n* = 5, meaning a cell produces 1 molecule for every 5 received plus 1. Note this is 2D and *n* is larger because the maximum amount of neighbours a cell can have is 4. Only the initial condition and stable state are shown. The gradient stabilises at timestep 32. C) Same as B but for a population size of 30 by 30 cells. The gradient stabilises at timestep 258.

The additive increase of +*a* in equation (11) rather than say a multiplicative increase prevents cells from increasing production rates to infinity, which would happen with a multiplicative increase if *a* was a constant. With the additive increase the gradient will eventually stabilise at a rate relative to the overall population size.

Stabilisation can still be achieved using multiplication if the rate of increase in production decreases as the actual rate of production increases, i.e. high rates of production will increase at lower rates compared to low rates of production which will increase at higher rates. For example, an equivalent multiplicative rule to the additive rule in equation (11) above would be as follows:

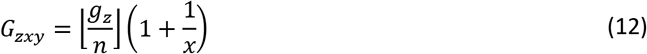

Where *x* is directly proportional to the value of molecules received. For example:

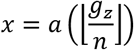

If *x* is directly proportional by a factor of 1 meaning in the above equation *a* = 1, then this is equivalent to saying *x* = amount of molecules received.

With this example, the 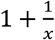 rate of increase is equivalent to adding *a* when *a* = 1 because *x* equals the amount of molecules received in the current timestep. For example if a cell receives 1 molecule it will produce 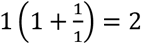. Or if a cell receives for example 8 molecules then it will produce 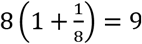. The absolute increase in value is constant and always 1, but the rate of increase relative to amount of molecules received decreases, which in the 2 examples went from an increase by a factor of 2 to a factor of 1.125.

**Figure 22 –.**
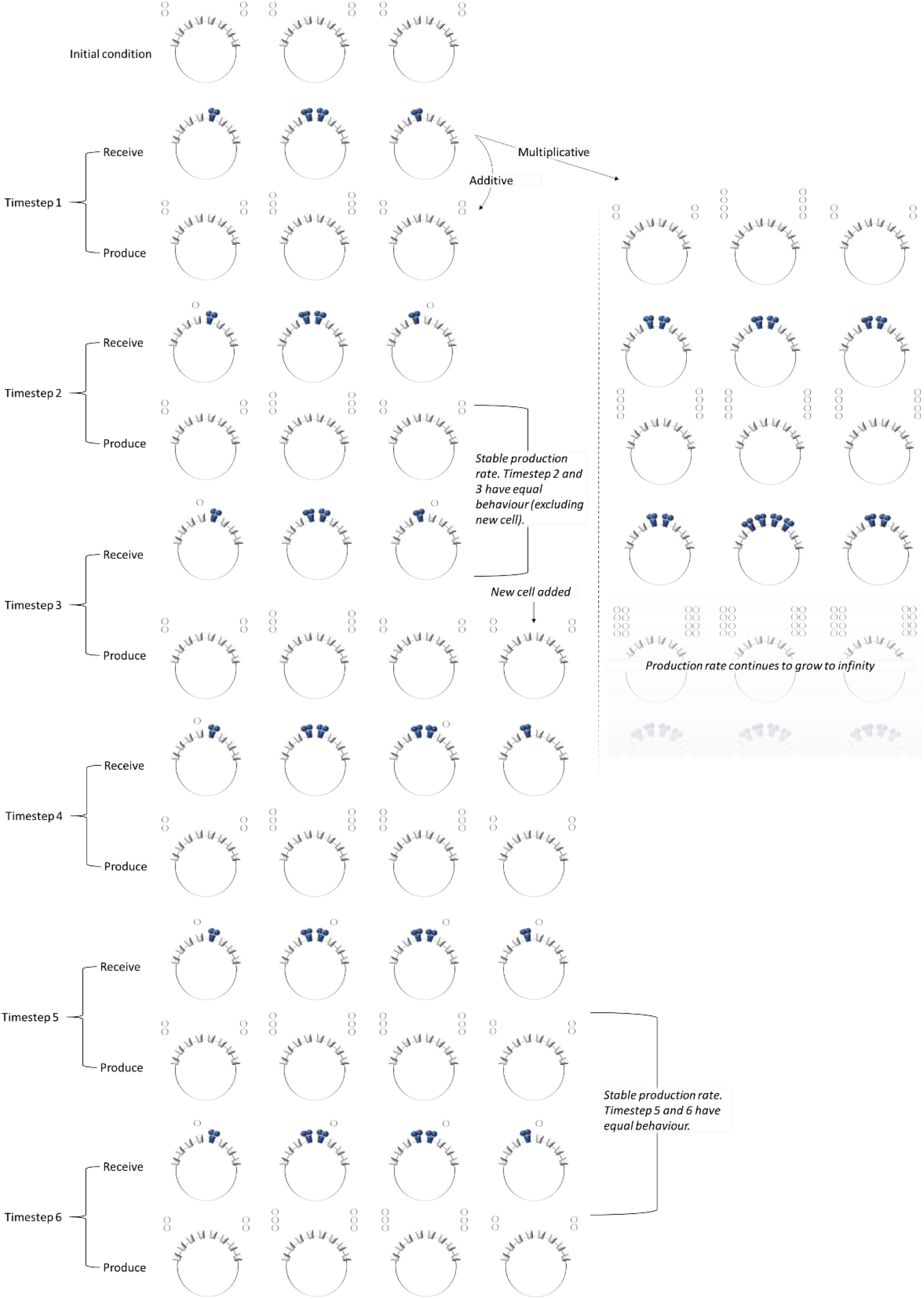
Conceptual diagram of gradient forming using equation (*11*) for the additive flow. Shows how the use of heterodimers enables a step threshold which corresponds to the amount of molecules a cell is receiving. The multiplicative flow uses a constant factor for *x* = 1 in equation (*12*), meaning the gradient does not stabilise because cells autoregulate by always multiplying the amount received by a constant of 2. In the additive flow, the gradient is stable at timestep 3 because it is repeat behaviour of timestep 2 for the 3 cells that already existed. At timestep 3 a new cell is also added and the gradient then stabilises again at timestep 6 which is repeat behaviour of timestep 5.

### Ligand inhibitors

In the previous example the production rate of the cells at the periphery and relative positions further into the population when signalling reaches a stable state increases with larger population sizes (see fig 20 and 21). This is because the rule used in equation (11) still allows a cell to receive a number of molecules from a single neighbour that is equal to or greater than an amount that would also be received when the cell is completely surrounded by neighbours which are producing at a lower rate. If one single neighbour can produce at a rate that causes a cell to increase its production rate when it is not completely surrounded by neighbours then this to some extent defeats the objective of the general rule described in the main text which is that cells increase their production rate only when all their neighbours are producing at a certain threshold rate.

Even though this occurs, it turns out that the gradient eventually stabilises, as shown in the previous section. However, the relative production rates of cells at the stable state increases significantly as population size increases, and it also takes significantly longer to reach the stable state as population size increases.

With use of an extracellular inhibitor it is possible to maintain consistent production rates of cells so that at the stable state the relative position of a cell will always result in the same production rate, regardless of overall population size (compare production rates in fig 23 and 24). The time taken to reach a stable state is also dramatically reduced. The way in which the inhibitor binds to gradient forming molecules and the rate at which cells produce an inhibitor can be set such that it causes increases in a single neighbours production rate to be ineffective on a cell unless all neighbours of that cell are producing at a rate greater than or equal to itself.

**Figure 23 –.**
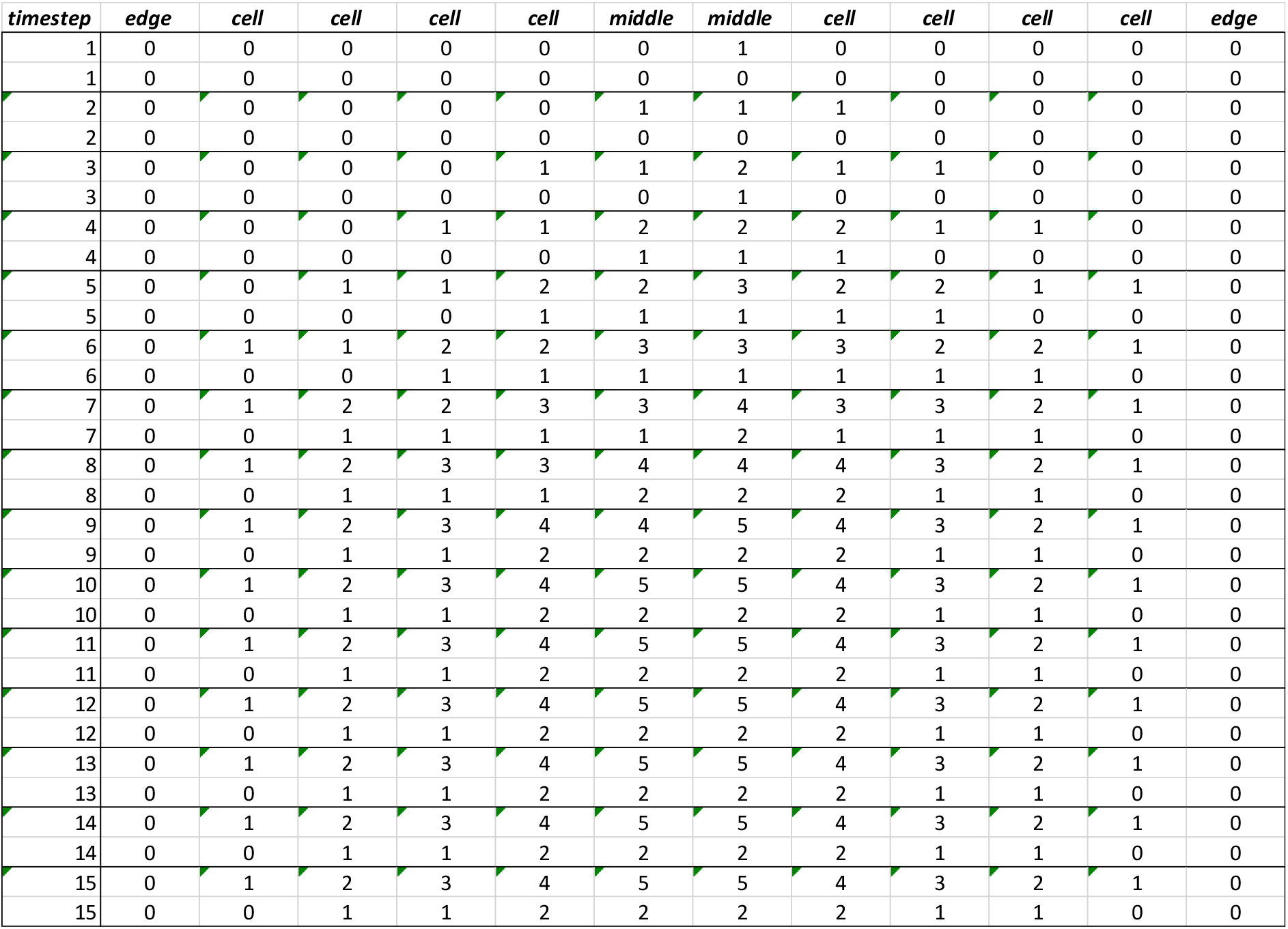
Table showing formation of a gradient using equation (*15*). There are 2 rows on each timestep, the top row represents the main gradient forming molecule and the second row is the amount of inhibitor produced as defined in equation (*14*). The gradient stabilises at timestep 10.

**Figure 24 –.**
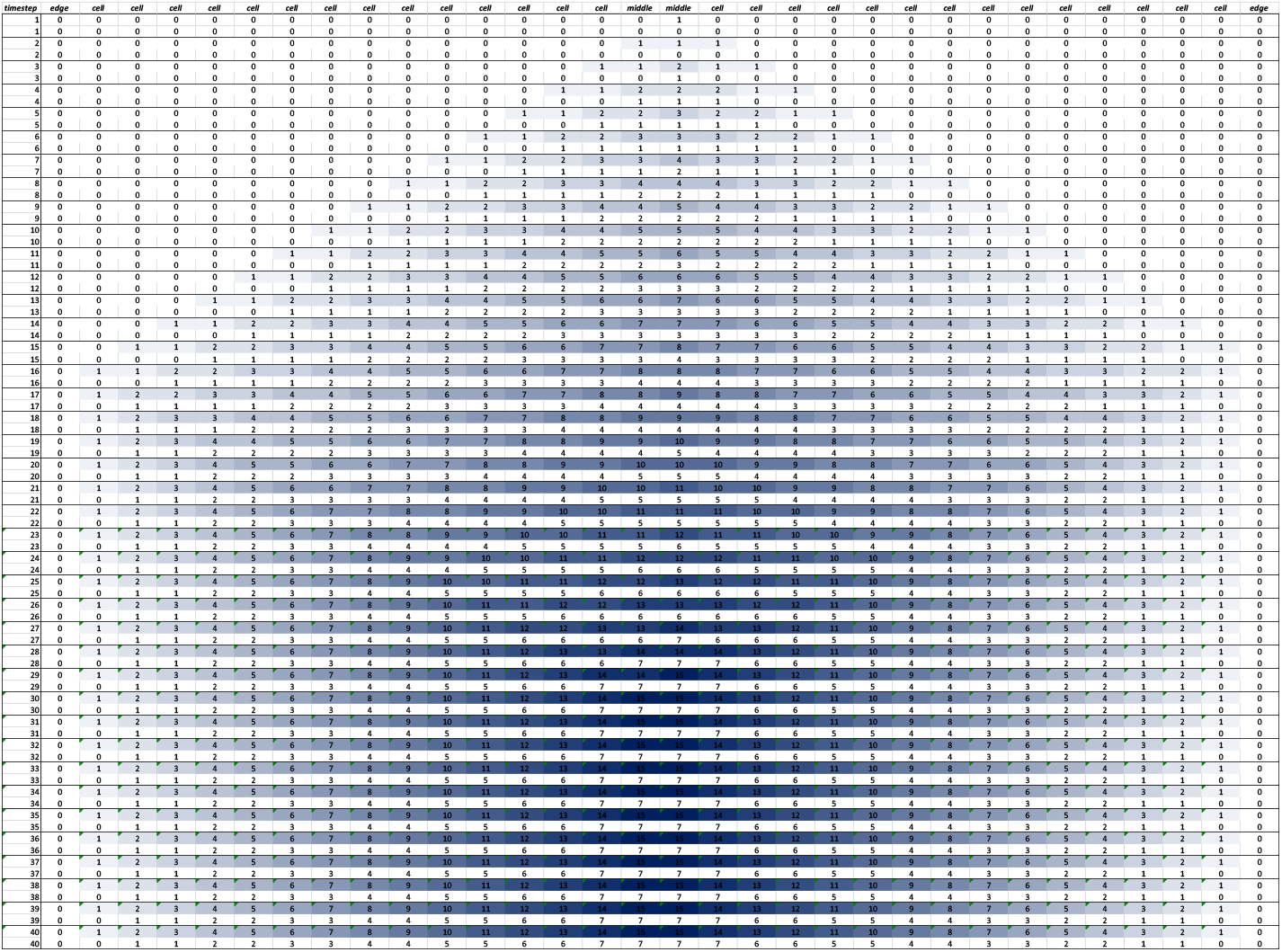
Table showing formation of a gradient using equation (*15*). There are 2 rows on each timestep, the top row represents the main gradient forming molecule and the second row is the amount of inhibitor produced as defined in equation (*14*). Cells showing the gradient forming molecule are shaded different strengths of blue to indicate the production rate of cells. The gradient stabilises at timestep 30.

In the 1D model for example, if the inhibitor is produced at a rate of half the gradient forming molecule, and the inhibitor binds itself in complexes of size *n* with higher affinity than binding to the gradient forming molecule, then when a cell and its neighbours produce at equal rates, the inhibitors stop binding to gradient forming molecules and allow cells to sense the additional molecule and increase their production rate.

The inhibitor production can be defined in a matrix *I* of size *trc* for 2D and *tr* for 1D. The production rate of inhibitor for each cell *C_zxy_ = I_zxy_* where:

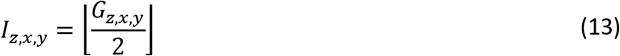

For 2D and each cell *C_zx_ = I_zx_* where:

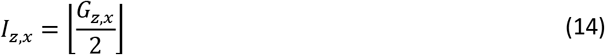

For 1D.

The autoregulation function for *G_zxy_* would be:

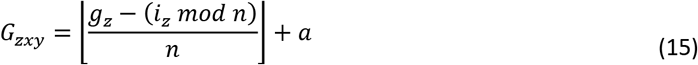

Were *i_z_* equals the sum of all inhibitor molecules produced by a cell and its neighbours in the previous timestep:

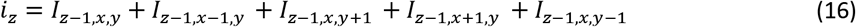

For 2D and

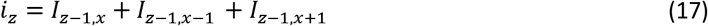

For 1D.

Equations (13) and (14) mean the rate of inhibitor production is equal to half the rate of gradient molecule production, rounded down to the nearest integer. The *i_z_ mod n* term in equation (15) encodes the behaviour that the inhibitor molecule has higher affinity to bind in complexes of size *n* with itself than the gradient forming molecule. The *g_z_* − (*i_z_ mod n*) term encodes the behaviour that leftover inhibitor molecules not in a complex bind one to one with gradient molecules in a dimer. When the inhibitor molecules bind in complexes of a size equal to the maximum number of a neighbours a cell can have plus 1, then when a cell is surrounded by neighbours producing at the same rate as itself, there will be no active inhibitors and this will allow the surrounded cell to receive a higher amount of molecule and increase its production rate. This means that *n* in the above equations should be equal to the maximum number of neighbours a cell can have plus 1.

Figure 23 shows the formation of a gradient in a row of 10 cells. There are two rows on each timestep, the first representing production value of the gradient forming molecule and second representing production value of the inhibitor molecules.

Figure 24 shows the formation of a gradient in a row of 30 cells. There are two rows on each timestep, the first representing production value of the gradient forming molecule and second representing production value of the inhibitor molecules.

This method of ligand inhibition will cause the gradient to grow in the exact same manner as the algorithm described in the main text, but only if the starting conditions are stable i.e. that cells begin by sending only 1 molecule to each of their neighbours. There is still a degree of fragility in this method of gradient formation because if for some reason a cell is producing at a higher rate than expected, it will cause the production rate of other cells in the population to stabilise at higher values. This is because cells are simply responding to the absolute value of molecules received and not able to determine the value of the lowest producing neighbour, subsequently this means there is not a strong enough method of downregulation when there is an overproducing cell.

### Receptor regulation

By regulating the amount of receptors on the cell surface, the issue explained at the end of the previous section on inhibitors can be resolved in certain cases due to a stronger level of downregulation when there are overproducing cells. The aim of regulating receptors is to cause production rates of cells to always converge to some value that corresponds exactly to a particular population size, regardless of initial production rates of cells, thereby creating almost equivalent behaviour and stability of the algorithm defined in equation (7) and (8) used in the main text. The algorithm used here works consistently for 1-dimensional rows of cells and for 2-dimensional populations up to a certain size. In certain cases with 2D, perturbations or random starting conditions can result in random signalling continuing indefinitely and a gradient never forming, this is shown in figure 28.

To limit the number of receptors on their surface, cells could simply upregulate the number of receptors if all of its existing receptors are being used and downregulate the number of receptors if not all receptors are being used. In addition to this, to make the gradient more stable, a rule can be used that causes unused receptors to inhibit or downregulate the production rate of the gradient forming signalling molecule.

Inhibition by receptors is conceptualised as occurring inside the cell, such that an unused receptor would somehow cause an inhibitory signal either through the process of degradation or by not being bound to any signalling molecule. A straightforward rule for inhibition is used here which is that each unused receptor blocks one activated receptor inside the cell. Receptors will be defined in a matrix *R* of size *trc*, where the *t* dimension represents each timestep and *r* and *c* represent the *x* and *y* axis of 2D space respectively. For the 1-dimensional model *R* is of size *tr*. Where the *t* dimension represents each timestep and *r* represents the row of cells.

At each timestep, a cell internalises all receptors and then replenishes a certain amount back to the cell surface. The values in each *R_zxy_* or *R_zx_* represents the amount of receptors replenished to the cell surface at each timestep.

This model incorporates the behaviours of the previous two sections which utilise ligands binding to receptors in complexes as well as the use of extracellular ligand inhibitors. Since there are now several factors at play, every single aspect of the signalling process is redefined here in piecemeal for clarity. See figure 25 for a schematic that corresponds to the equations below.

**Figure 25 –.**
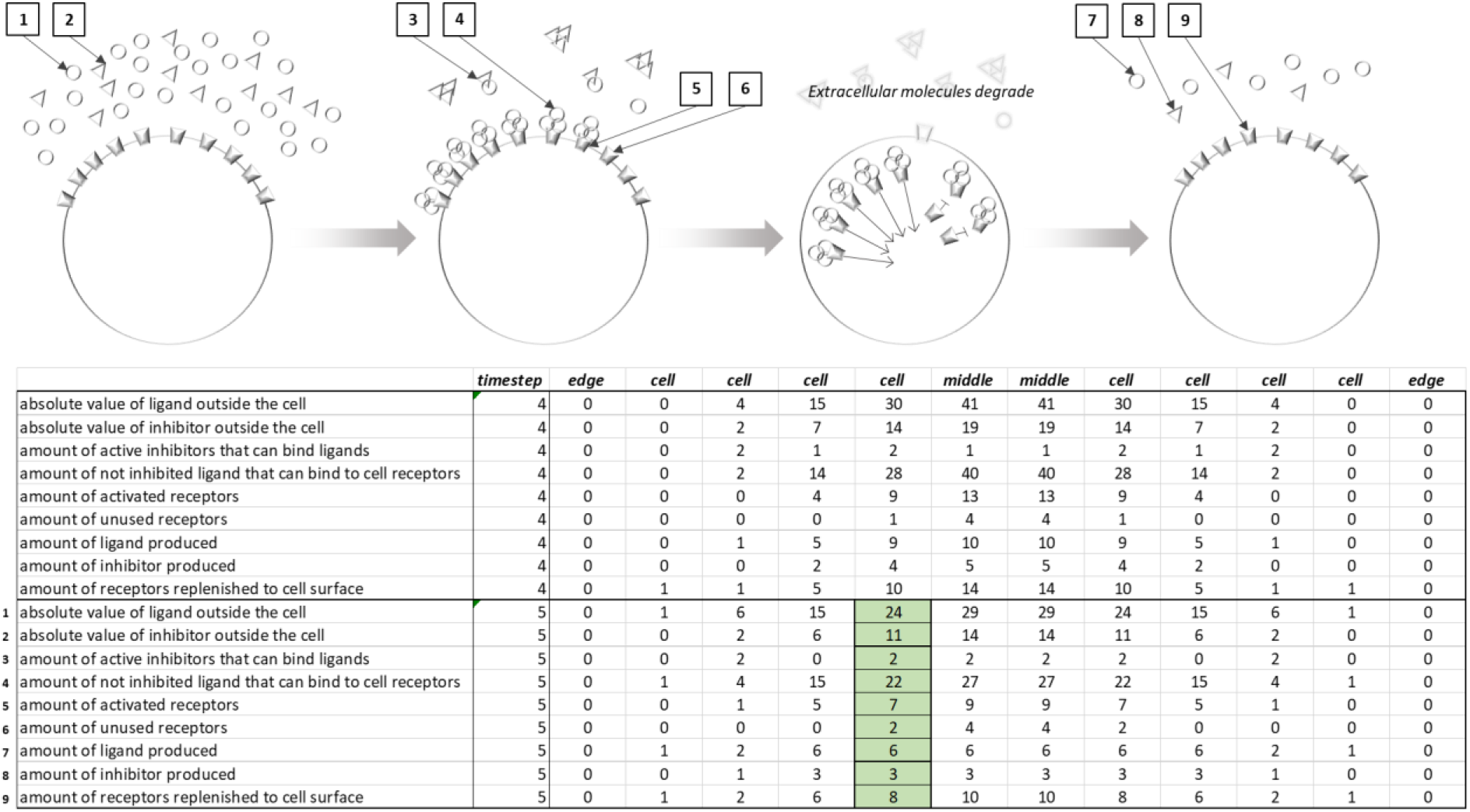
Schematic of the regulatory activities of a cell using all equations defined in this section. The diagram corresponds to the values in the highlighted cells in timestep 5 in the table. The table is an extract of a gradient forming in a row of 10 cells with initial conditions that the middle cells had a production value of 30 ligands, which is too high for a population of this size (central cells should stabilise with a value of 5, see figure 11), therefore the cells are in the process of downregulating in order to converge to correct and stable production rates. It can be seen that the total value of ligands and inhibitors the cell received from itself and neighbours is 24 ligands and 11 inhibitors. Subsequently the inhibitors bind with higher affinity into trimers and any remaining bind 1 to 1 with ligands. As there are 11 inhibitors, this results in 3 trimers and 2 binding to one ligand each. Uninhibited ligands are then able to bind to receptors in trimers. As there are 22 uninhibited ligands, this means 7 trimers can be formed to bind to and activate 7 receptors, with one ligand remaining that cannot bind to receptors or inhibitor. The receptors are then internalised and 2 of the 3 unused receptors inhibit 2 of the activated receptors, meaning only 5 of the 7 activated receptors can transduce signals to the nucleus. The cell then produces 6 ligands as per equation (*20*). It also produces 3 inhibitors as per equation (*13*) and replenishes 8 receptors as per equation (*21*). Note that although the diagram shows 6 ligands and 3 inhibitors produced, this is the amount the cell sends to each of its neighbours and itself. Since it has 2 neighbours the absolute value produced will be 3 times this amount which is 18 ligands and 9 inhibitors.

At each timestep ligands will be sent to a cell from each of its neighbours plus itself. Therefore, the *absolute value of ligand outside the cell* is equal to the sum of that produced by the cell and all of its neighbours in the previous timestep, which is represented by *g_z_* as already defined in equation (5):

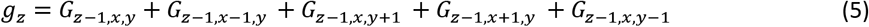

A similar rule applies for the *absolute value of inhibitor outside the cell* which is equal to the sum of inhibitors produced by the cell and all of its neighbours in the previous timestep, which is represented by *i_z_* as already defined in equation (16):

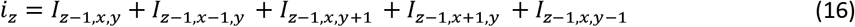

As mentioned in the previous section, inhibitors have an affinity to bind themselves in complexes of size *n* and any remaining will bind to ligands in dimers. Therefore, the *amount of active inhibitors that can bind ligands* is:

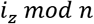

Where *n* = maximum number of neighbours a cell can have.

The *amount of not inhibited ligand that can bind to cell receptors* is the absolute value of ligand outside the cell minus the amount of inhibitors that can bind ligands:

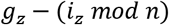

The *amount of activated receptors a_z_* is the amount of complexes of size *n* that can be formed by not inhibited ligand:

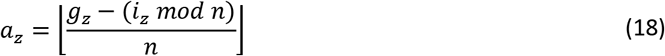

The *amount of unused receptors u_z_* is the amount of receptors on the cell surface that were not activated by ligands, which is equal to the amount of receptors replenished to the cell surface in the previous timestep *R*_*z*−1,*x,y*_ (and therefore the amount available to sense ligand in the current timestep), minus the amount of activated receptors. The minus 1 accounts for a ‘spare’ receptor that gets produced to enable cells to sense increases in ligand and subsequently upregulate production. This additional receptor should not cause inhibition, otherwise the rate of inhibition will be too great and the gradient will not grow:

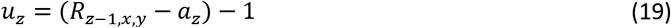

The *amount of ligand produced* is the amount of activated receptors minus the amount of unused receptors, plus 1. The plus 1 is the rate of upregulation:

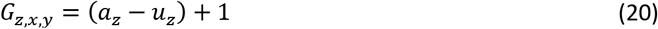

The *amount of inhibitor produced* is the same as that defined in equation (13) in the previous section which is relative to the amount of ligand produced:

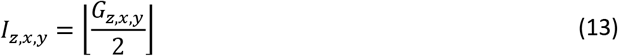

The *amount of receptors replenished to cell surface* is the amount of activated receptors plus 1. The plus 1 accounts for the ‘spare’ receptor which is there to sense additional ligand and enable upregulation when extracellular ligand increases:

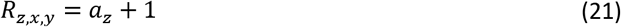

Figure 26 shows formation of a gradient using the above equations in a row of 10 cells.

**Figure 26 –.**
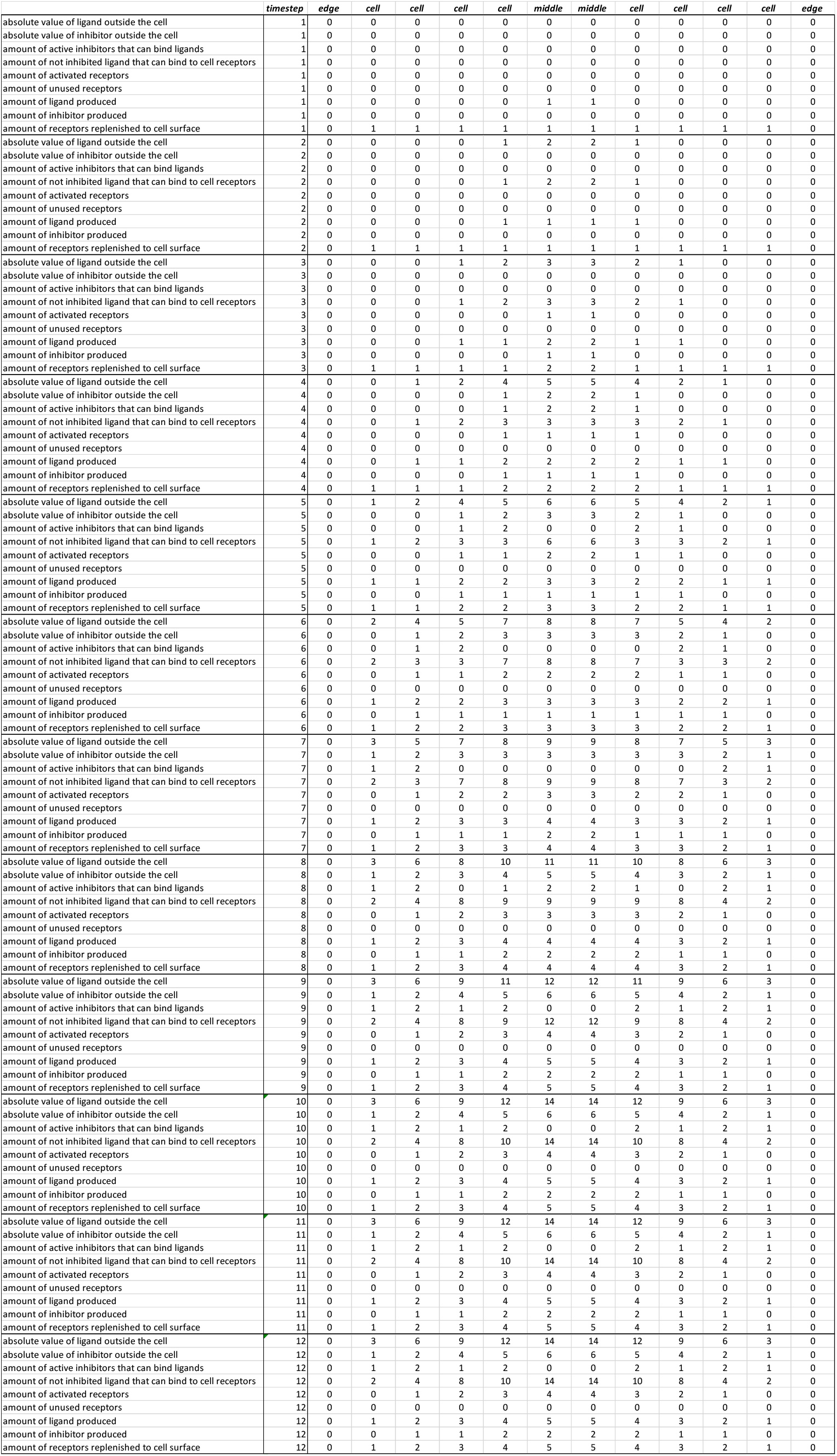
Table showing the formation of a gradient over time using the regulatory rules defined in all equations in this section, where *n* = 3. Each timestep has a row to represent the value of each related equation for that cell. The gradient stabilises at timestep 9, where the production of ligand will be the same for all cells in subsequent timesteps.

Figure 27 shows formation of a gradient from random starting conditions in a 10 by 10 grid of cells.

**Figure 27 –.**
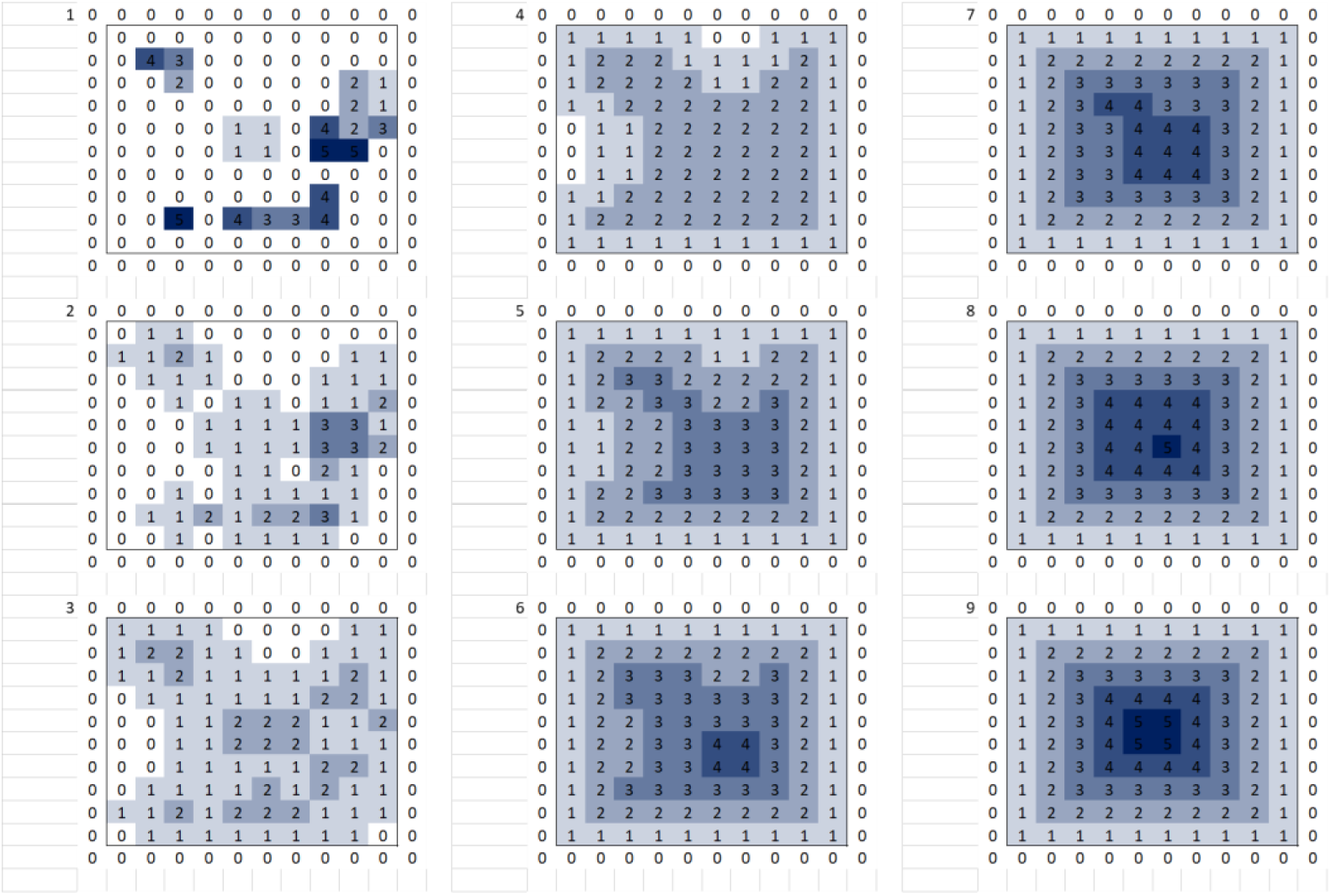
Formation of a gradient over time with uneven initial conditions in a 10 by 10 grid of cells using the regulatory rules defined in all equations in this section, where *n* = 5. Cells are shaded different strengths of blue to indicate the production rate. Moving from top to bottom then left to right, each image shows the production rates of cells at each timestep. 9 timesteps are shown. The gradient stabilises at timestep 9, meaning cells exhibit the exact same signalling behaviour in subsequent timesteps.

Figure 28 shows formation of a gradient in a 30 by 30 grid of cells. Certain starting conditions in a population of this size actually result in random signalling behaviour from cells and no gradient formed. This is likely due to the receptor regulation algorithm being too simplistic and simply subtracting very large values to regulate production when there are overproducing cells. This appears to result in cells downregulating too heavily and constantly ‘resetting’ from zero, this behaviour cascades throughout the population but never finds equilibrium.

**Figure 28 –.**
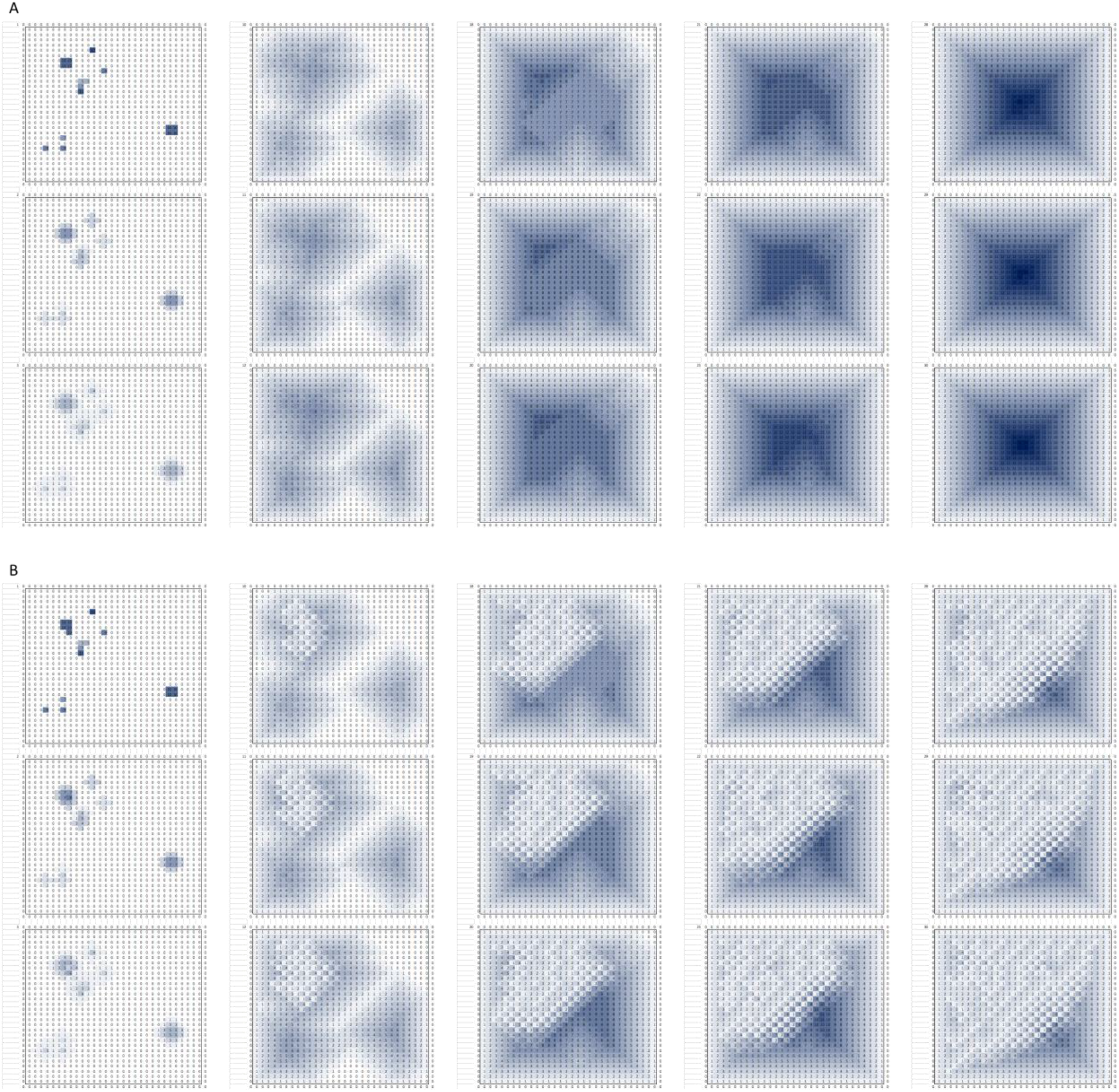
Formation of a gradient over time with uneven initial conditions in a 30 by 30 grid of cells using the regulatory rules defined in all equations in this section, where *n* = 5. Cells are shaded different strengths of blue to indicate the production rate. Moving from top to bottom then left to right, each image shows the production rates of cells at each timestep. In A and B the same timesteps are shown up to timestep 30. Some timesteps are not shown to save space. In A) it can be seen that the gradient stabilises at timestep 30. In B) there is only one cell different in the initial conditions yet the gradient does not stabilise, all subsequent timesteps look similar to timestep 30, with random low values of signalling occurring throughout the population.

## Appendix 2 – Scale invariance of opposing gradients

Scale invariant opposing gradients is taken to mean that there are 2 gradients of different molecule types generated in a population of cells and the relative size of these 2 gradients always remains in the same proportion no matter what the overall size of the population is. For example if the ratio is 2:3 then in a row of 5 cells, 2 cells incorporate one gradient and 3 cells incorporate the other, if this was in a row of 10 cells then it would be 4 and 6.

The autoregulation rule in equation (8) is used to generate gradients and any signalling rules used in addition to this are described below. This section is to show opposing gradients that scale according to population size can be formed using the model defined in this paper and not making any claim that real biological systems use the same or similar mechanisms. The scale invariance model is discussed only in the context of the 1-dimensional model and has only been implemented for the 1-dimensional model.

**Figure 29 –.**
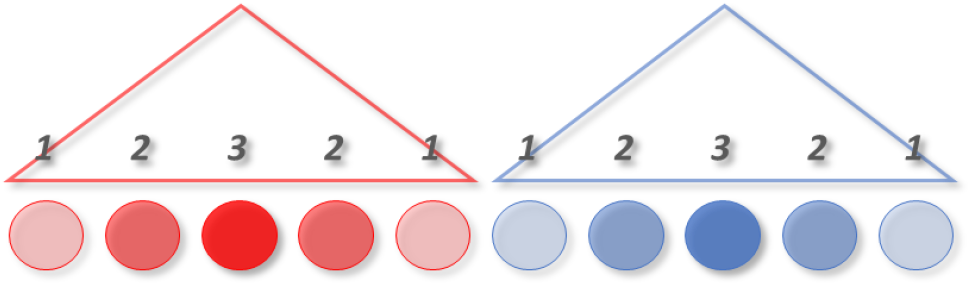
Schematic diagram of opposing gradients in a row of 10 cells. The gradients are in a ratio of 1:1.

In short, scale invariance is achieved by cells in each of the two gradients producing a secondary signal in addition to the gradient forming signal, which acts as a direct readout of gradient size for each cell in the population. This is a flat, consistent signal that spreads across all cells and its size is dictated by the production rate of the centre cells of the gradient, which are those that are producing the highest amount of gradient molecule and have accurate information about the overall population size for the gradient they are part of. Cells at the boundary of the two gradients get information about the size of both gradients and are therefore able to modify their signalling behaviour to take the two gradients towards the desired ratio.

### Information required for scale invariance

In the context of this model, the gradient signal alone is not enough information for a cell to accurately determine its position within that gradient relative to other cells, because the same absolute number of signalling molecules that a cell receives could mean for example that it is near the centre of a small population or near the edge of a larger population (fig 30).

**Figure 30 –.**
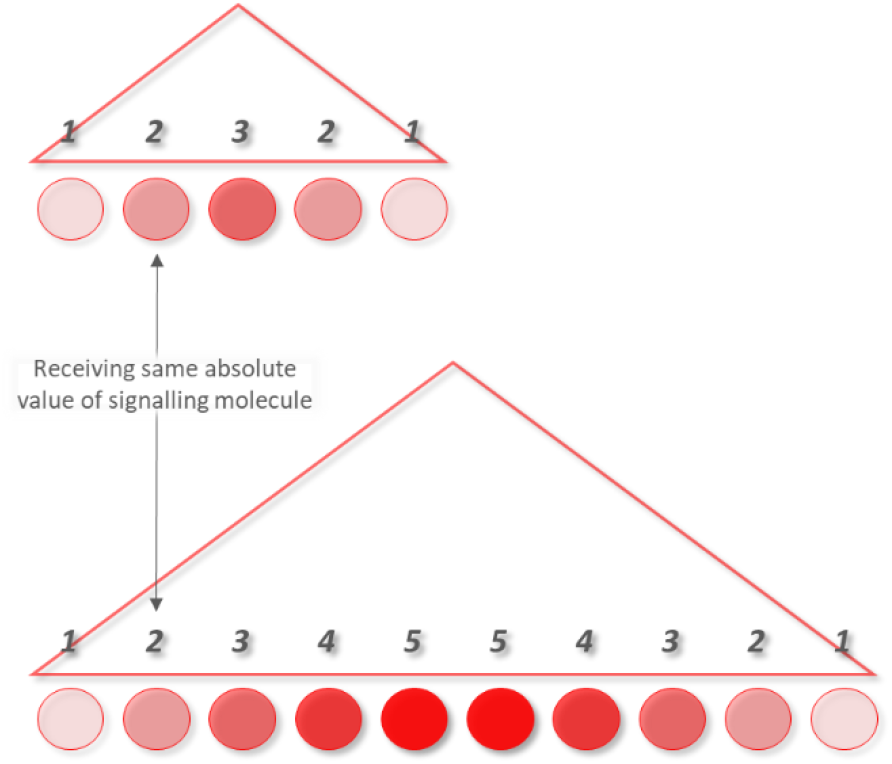
Cells will receive the same absolute value of signalling molecule according to their distance from the edge of the population, no matter how large the overall population size is.

It would be possible for a cell to determine its relative position in a gradient if it had information about the overall population size, in addition to the positional information provided by the gradient forming molecule/signal. With these two pieces of information, any cell could determine its relative position by comparing the absolute value of gradient molecule which represents position, against the value of population size.

As shown in the main text, the central cell in a population has accurate information about population size. If a signal could be propagated to all cells that represents the central cells calculation of population size, then all cells can have accurate information about their relative position within a gradient (fig 31).

**Figure 31 –.**
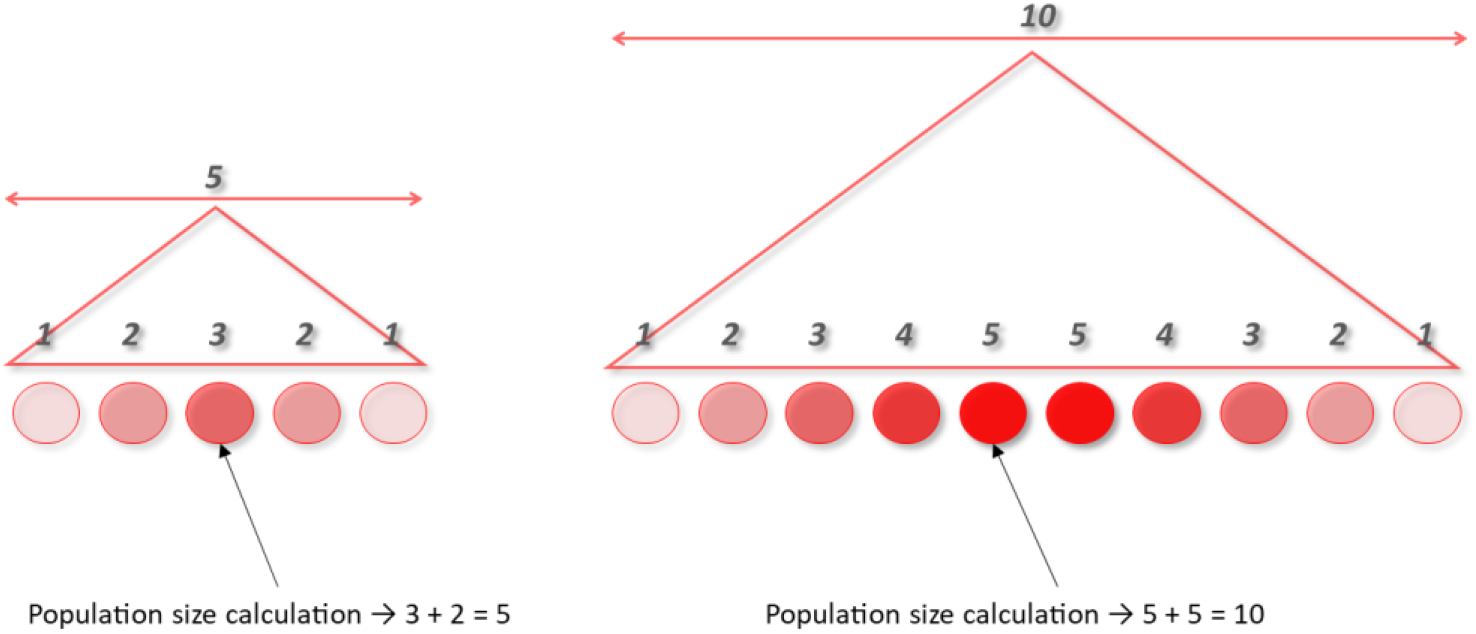
Central cells can accurately determine the population size by adding the value of molecules they are producing to the value produced by their highest producing neighbour.

Using this system, it is possible to combine two opposing gradients in a population of cells such that they scale according to overall population size. This works on the basis that cells at the boundary of the two gradients will receive signals from both gradients and therefore have information about the population size of each gradient, subsequently allowing them to adjust their signalling behaviour so that the gradients move towards the desired ratio (fig 32).

**Figure 32 –.**
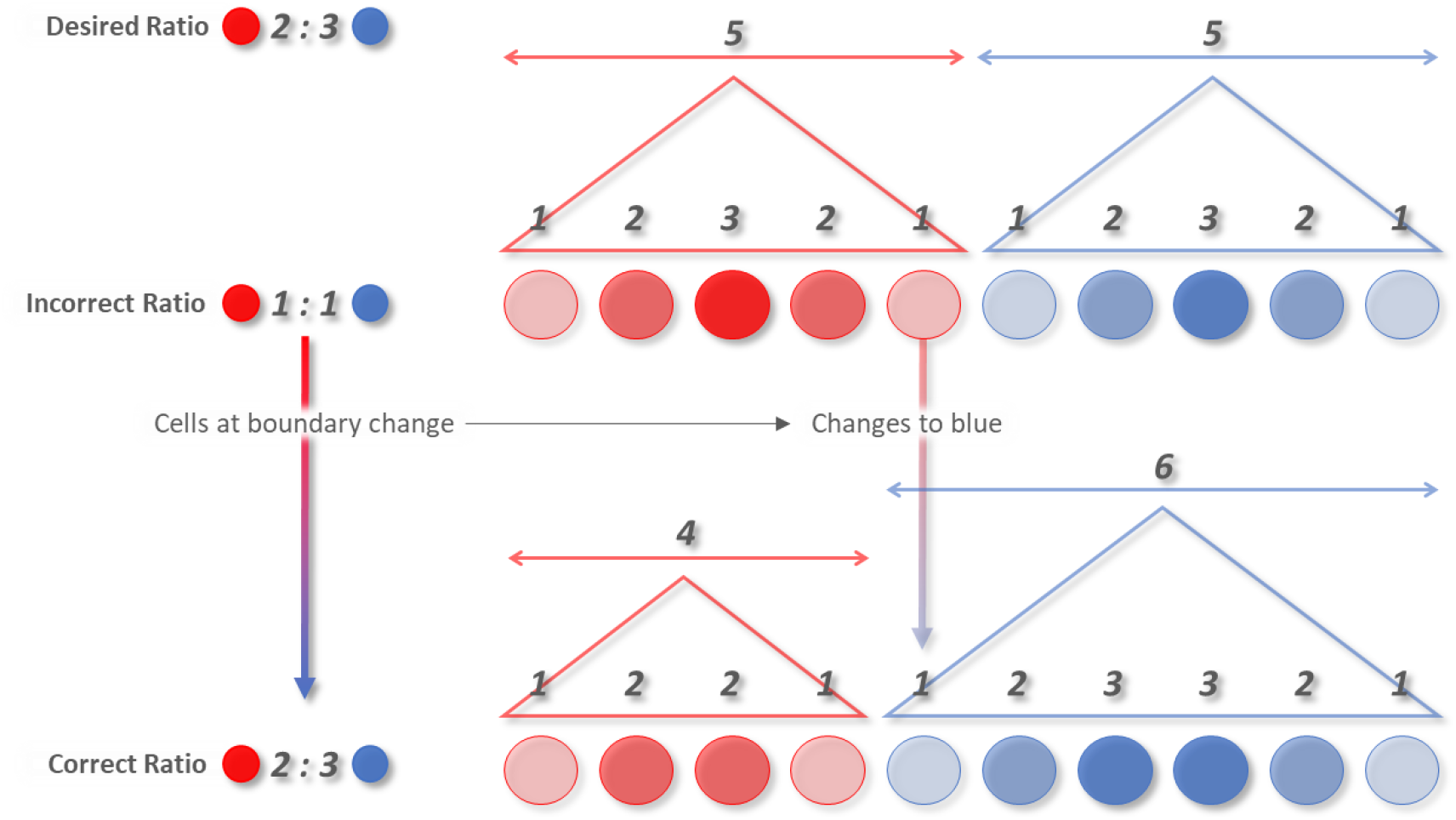
Boundary cells receive information about the size of both gradients and can adjust their signalling to cause the overall population move towards the desired ratio.

The next two sections describe how the population size signal can be calculated and propagated to all cells in a gradient, and also how boundary cells adjust their signalling behaviour so that the gradients always move towards the desired ratio.

### Population size signal

The central/highest producing cell has accurate information about the size of the population and can therefore propagate this information all the way to cells at the edge of the gradient by sending out an additional signal which is a calculation of the population size. This ‘population size’ signal will be passed on cell by cell until it reaches the edge of the population and acts as a direct readout of population size for every cell.

How does a cell calculate the population size? Since each cell is receiving a certain amount of gradient signalling molecules from its neighbours and producing a certain amount itself, these values can be taken as a number and used to calculate the actual population size. The following calculation turns out to give an accurate population size:

> *population size = amount of molecules a cell is producing + amount of molecules being received from its highest producing neighbour*

The result of this equation when calculated by the central cell will be an accurate calculation of population size. Other cells however will not produce an accurate calculation of population size using this equation (fig 31).

If every cell is using the same rule set, then every cell will be able to calculate its estimate of the population size and pass this onto its neighbours. This is undesirable because only the centre cell is making an accurate calculation of population size and we want this signal to be passed around the population, not the inaccurate calculation made by other cells. This means each cell needs to decide whether to make its own calculation of population size and send this to its neighbours, or simply pass on the population size signal received from its neighbours.

This can be accomplished by saying that if a cell is producing a gradient signal greater than or equal to all of its neighbours, then it should make its own population size calculation and send that signal to each of its neighbours. Else if any neighbour is producing more than it, then take the population size signal of the highest producing neighbour and pass that on to all neighbours.

This subsequently guarantees that a flat signal will be propagated across the population that is equal to the population size calculation made by the central cell (which is a correct calculation).

### Signalling behaviour of boundary cells

All cells in the overall population have the exact same rule set, so there needs to be a reason why each gradient doesn’t expand into the other. This could be explained by saying that each molecule inhibits the other, or that a cell can only make a binary decision and produce one type of molecule or the other.

Gradients in real biological systems often merge into each other at the boundary and it is conceivable that a cell could respond to and produce both types of morphogen/signalling molecule, including in this model. It turns out to be easier for the purposes of demonstrating this scale invariance concept to take the stance that cells will only produce one type of molecule or the other, therefore the binary choice method is used for this model whereby a cell can only produce one type of molecule or the other, but it can receive/sense both types of molecule at all times.

If the opposing gradients are not in the desired ratio, there needs to be a way in which cells will change which signalling molecule they produce so that as a population they move towards the desired ratio. It makes sense for this change to occur at the boundary of the two gradients, meaning that if a cell at the boundary changes which signalling molecule it is producing, then one gradient becomes one cell larger and the other one cell smaller. Cells not receiving both signalling molecules should never change because they will not be at the boundary of the gradients (fig 32).

This means a rule is needed that causes cells to decide which signalling molecule to produce if they are receiving both types. This decision should of course be based on moving towards the desired ratio between the two gradients. Since cells at the boundary will be receiving the population size signal for both gradients, it is a simple calculation of the two population readouts, if the ratio is too high or low one way or the other then the cell will decide to become part of the population that takes the overall gradients towards the desired ratio.

### Synchronicity

The model described so far may appear to make it possible to construct two opposing gradients in a population of cells. However, for quite interesting reasons the model as described up to this point will not work. There is still a subtle problem which is related to allowing changes in signalling in one part of the population to propagate to all cells in the rest of the population.

When a cell decides to change which signal it is producing, it must do so with a certain degree of confidence that it is making the correct decision. If a cell changes from one gradient to the other, then the production rates of cells further into the population will not change immediately due to the fact that the sudden change in signalling at the boundary will only impact the immediate neighbours of boundary cells. In subsequent timesteps, the neighbours of neighbours of boundary cells will be impacted, and then neighbours of neighbours of neighbours and so on at each individual timestep until the change in signalling has impacted cells all the way to the centre of the population. This means that in order for a change at the boundary to impact the overall behaviour of the entire population, the change in signalling needs to propagate all the way down to the central cell, which is also where the correct population readout is coming from. This will cause a change in population size signal being sent from the central cell, which then needs to propagate back through the population and out to the boundary cells so that they have a correct readout of population size. In summary, when an edge cell changes its behaviour, it causes a wave of changes in signal production rates which propagate from the population edge to the centre and back out to the edge again. Once the changes have stabilised, the edge cells will have correct information about the gradients. Before this point they cannot be certain that they have correct information because they could still be receiving signals that are a result of cells in other parts of the population that have not yet been impacted by the change of signalling at the boundary.

In order to prevent cells from changing behaviour with inaccurate information there needs to be some mechanism that inhibits cells from making any changes until they are confident that signals they are receiving are stable and reliable.

This can be done by using a variation of the population size rule already defined. The difference will be that instead of immediately matching a neighbouring cells population size signal value, a cell will take a certain amount of time to move towards that value. Importantly, the amount of time a cell will take to get to that value is relative to the overall population size and that cells position in that population.

In this model the population size molecule production rate can simply be increased by 1 if the cell is producing less gradient molecule than its neighbour. This means that if a cell decides it needs to mimic its neighbours population production rate rather than calculate its own, it moves gradually towards that value rather than immediately changing to match its neighbours production rate. If the rate of change of production of moving towards its neighbours production value is set correctly, it can be such that it will always allow time for the gradient value to propagate throughout the population before the population value has reached a stable state.

A cell can decide whether its population value is at a stable state based on whether it is producing the same amount as its neighbours. If either of its neighbours are producing a different population value, then it is not stable, if they are all producing the same population value, then it is stable.

Using the above method, we can then say that when a cell changes from producing one molecule to another, or if it suddenly starts receiving both molecule types, then they ‘reset’ their population value to 1. This acts in an inhibitory manner because it must count up again to the actual population size which gives time for the wave of changing of new gradient signals to propagate throughout the population. This ensures that when a cell changes, the one that changed resets its value and the cell next to it in the opposing gradient (which has gone from being inside the population to being at the edge) also resets its value. The cells then ‘wait’ for a stable state to occur before any change can be induced again.

### Implemented examples

The figures below (33, 34, 35, 36) show various implementations in Microsoft Excel using the signalling rules described above.

**Figure 33 –.**
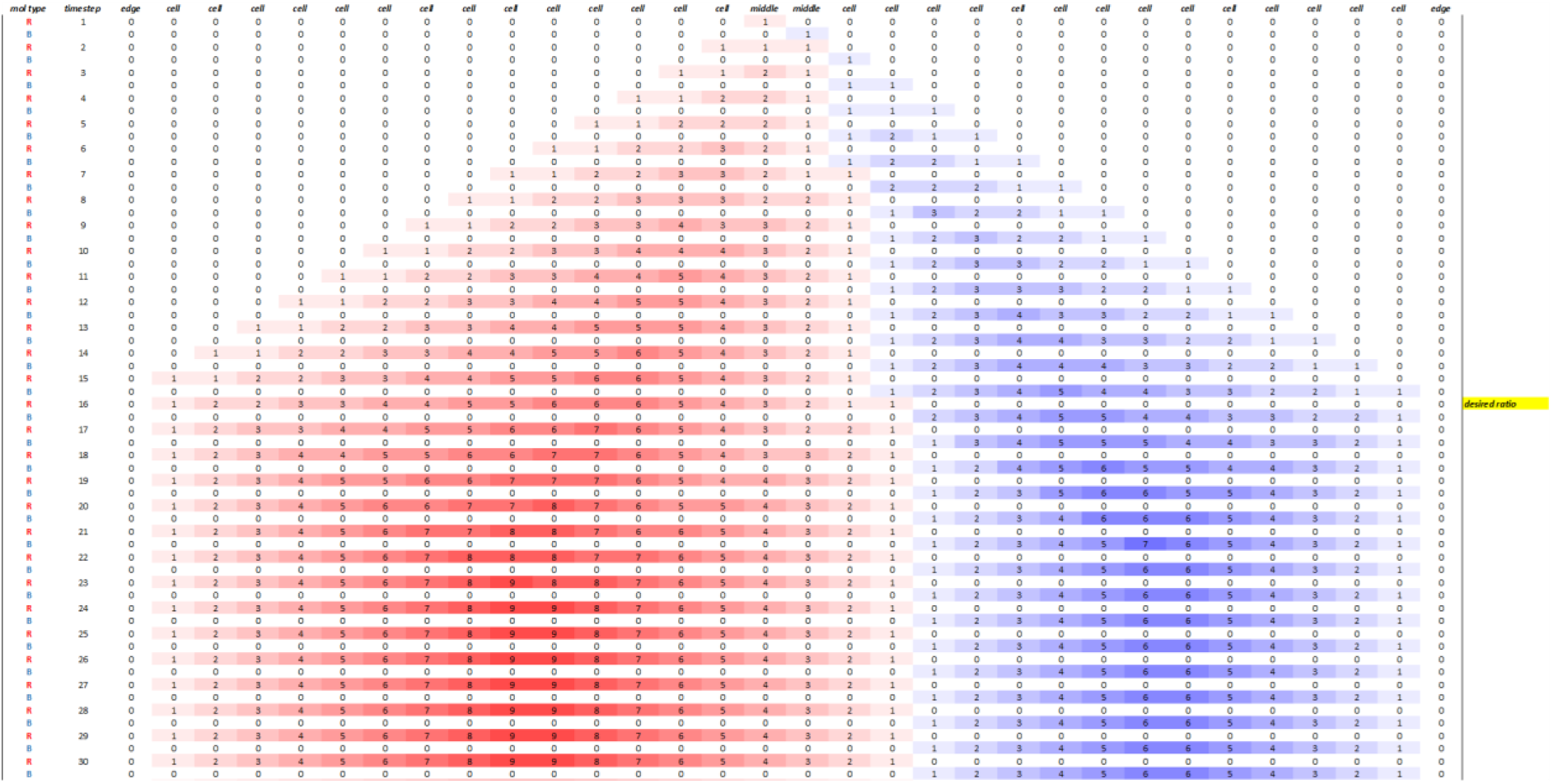
Microsoft Excel implementation of two opposing gradients that scale according to population size in a row of 30 cells. There are two rows per timestep, one each to represent the value produced by a cell for each molecule type ‘R’ and ‘B’. The yellow marker indicates the point at which the desired ratio is reached. The desired ratio of R:B in this example is 3:2.

**Figure 34 –.**
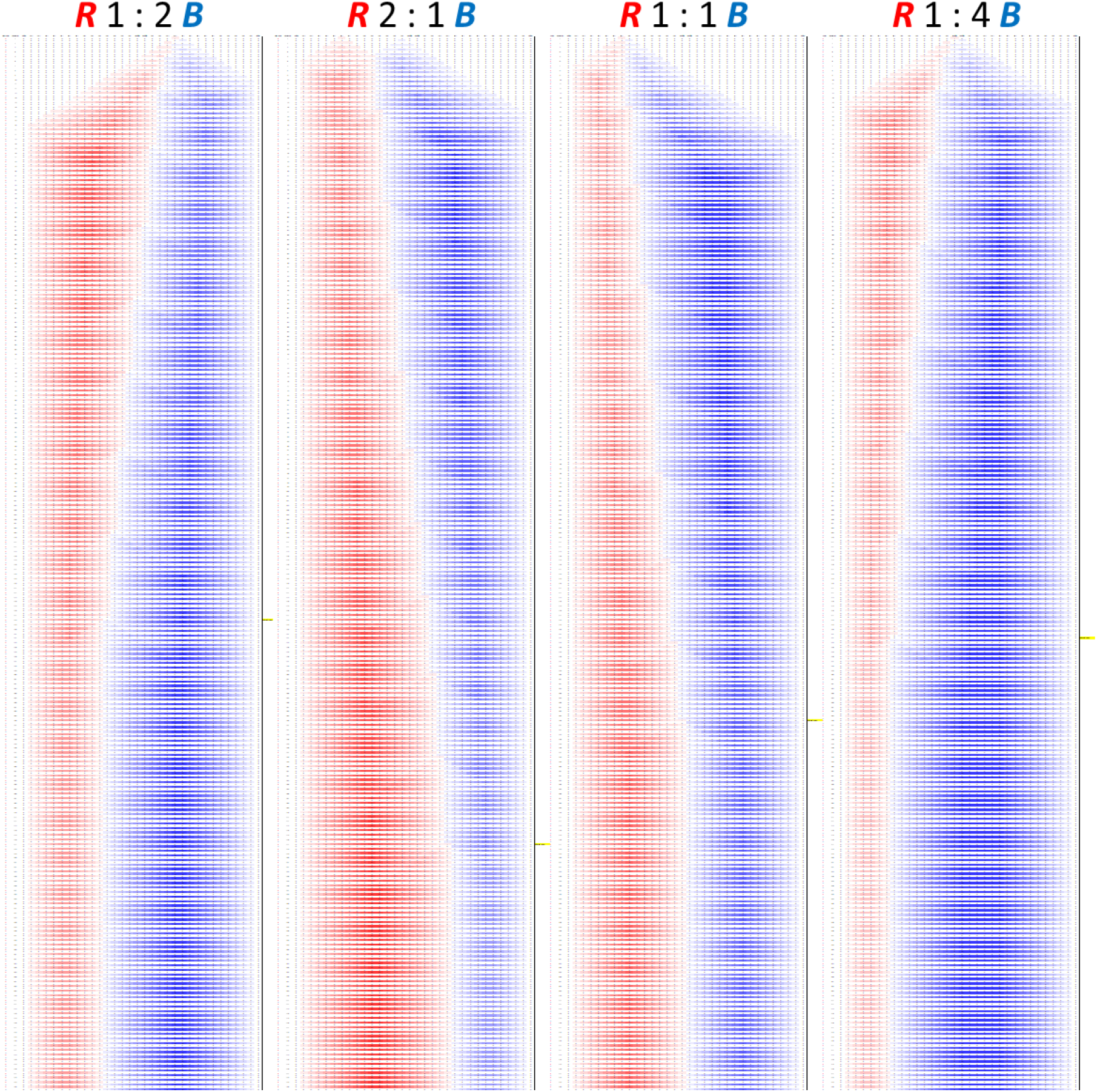
Various examples of Microsoft Excel implementations of two opposing gradients that scale according to population size in a row of 30 cells. There are two rows per timestep, one each to represent the value produced by a cell for each molecule type ‘R’ and ‘B’. The yellow marker on the right hand side of each example indicates the point at which the desired ratio is reached. The desired ratios are shown at the top of each example. The initial conditions vary for each implementation to show that the gradients will stabilise from various initial conditions, so long as there are not overlapping cells that produce different molecules of R and B.

**Figure 35 –.**
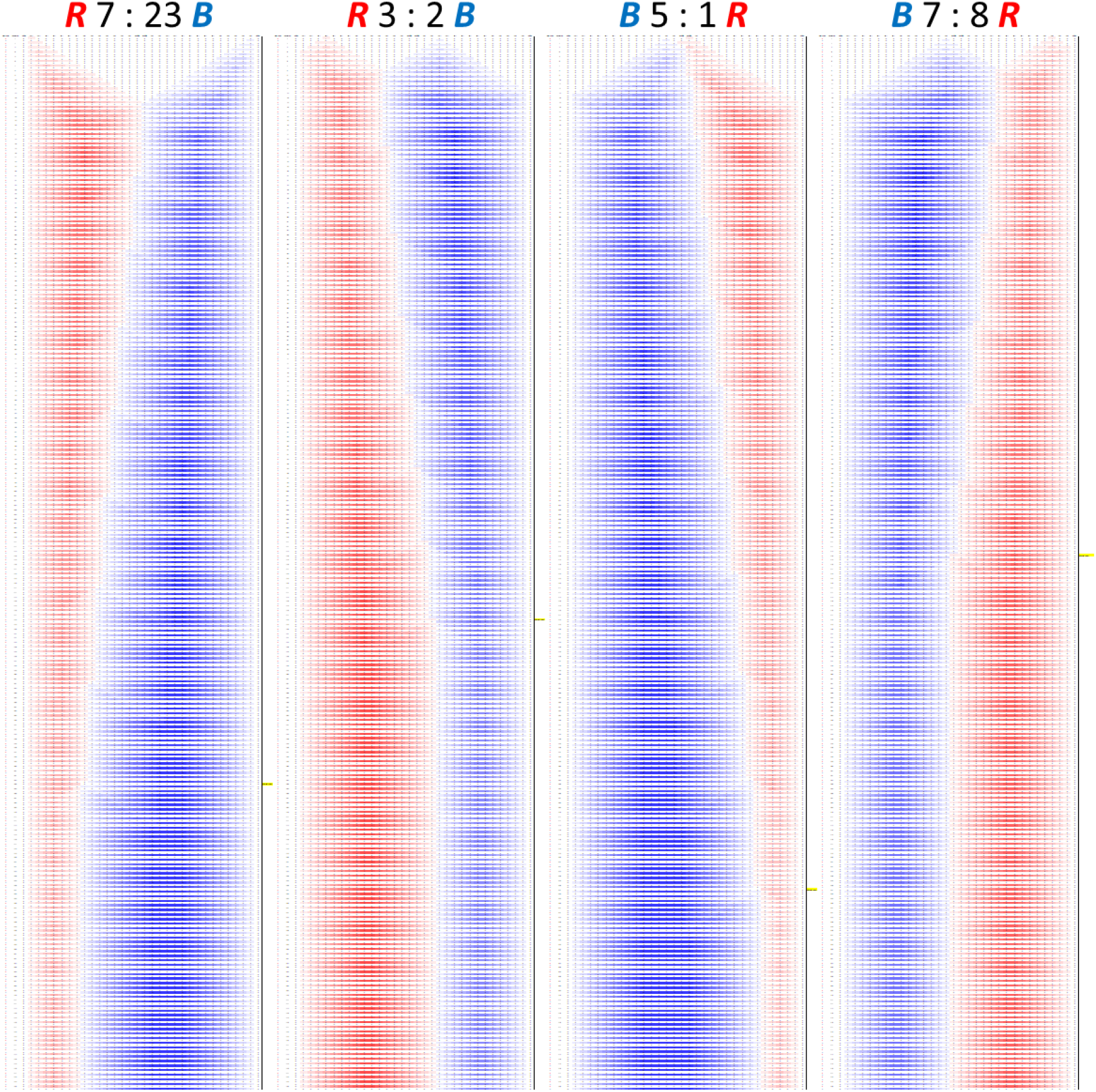
Various examples of Microsoft Excel implementations of two opposing gradients that scale according to population size in a row of 30 cells. There are two rows per timestep, one each to represent the value produced by a cell for each molecule type ‘R’ and ‘B’. The yellow marker on the right hand side of each example indicates the point at which the desired ratio is reached. The desired ratios are shown at the top of each example. The initial conditions vary for each implementation to show that the gradients will stabilise from various initial conditions, so long as there are not overlapping cells that produce different molecules of R and B.

**Figure 36 –.**
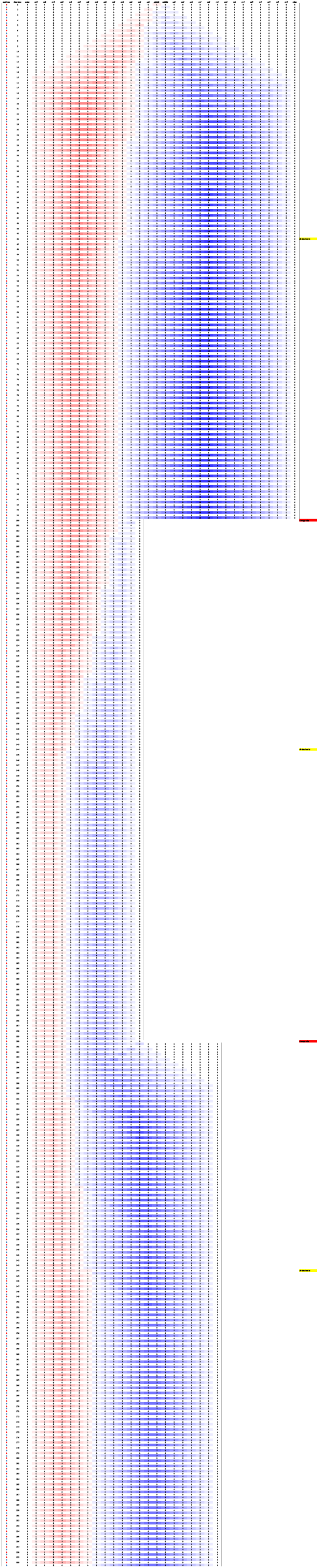
Microsoft Excel implementation of two opposing gradients that scale according to population size where the size of the population is changed at two points during the development of the gradient. There are two rows per timestep, one each to represent the value produced by a cell for each molecule type ‘R’ and ‘B’. The yellow marker on the right hand side indicates the point at which the desired ratio is reached, the red markers indicate the point at which the population size is changed. The desired ratio of R:B is 1:2. The population size starts of as a row of 30 cells. The desired ratio is reached at timestep 46 and the population size is then reduced to 12 cells at timestep 100. The two gradients then move towards the desired ratio again which is reached at timestep 144. At timestep 201 the population size is increased to 15 cells and once again the gradients move towards the desired ratio which is reached at timestep 244.

In summary, cells are performing the following general functions:

- Producing a gradient forming signal.
- Will only produce a maximum of one type of gradient forming signal at any point in time.
- Producing a population size signal related to the gradient forming signal they are producing.
- If receiving both types of signal, will choose to produce the signal that takes the two gradients towards the desired ratio.
- Will only change signalling behaviour if population size signals are stable, which is when they are producing the population size signal at the same rate as their immediate neighbours.

